# A Dendritic Disinhibitory Circuit Mechanism for Pathway-Specific Gating

**DOI:** 10.1101/041673

**Authors:** Guangyu Robert Yang, John D. Murray, Xiao-Jing Wang

## Abstract

In this work we propose that a disinhibitory circuit motif, which recently gained experimental support, can instantiate flexible routing of information flow along selective pathways in a complex system of cortical areas according to behavioral demands (pathway-specific gating). We developed a network model of pyramidal neurons and three classes of interneurons, with connection probabilities constrained by data. If distinct input pathways cluster on separate dendritic branches of pyramidal neurons, then a pathway can be gated-on by disinhibiting targeted dendrites. We show that this branch-specific disinhibition can be achieved despite dense in-terneuronal connectivity, even under the assumption of random connections. We found clustering of input pathways on dendrites can emerge through synaptic plasticity regulated by disinhibition. This gating mechanism in a neural circuit is further demonstrated by performing a context-dependent decision-making task. Our findings suggest a microcircuit architecture that harnesses dendritic computation and diverse inhibitory neuron types to subserve cognitive flexibility.

## Introduction

Distinct classes of inhibitory interneurons form cell-type specific connections among themselves and with pyramidal neurons in the cortex^1,2^. Interneurons expressing parvalbumin (PV) specifically target the perisomatic area of pyramidal neurons. Interneurons expressing somato-statin (SOM) specifically target thin basal and apical tuft dendrites of pyramidal neurons^3^. In-terneurons expressing vasoactive intestinal peptide (VIP) avoid pyramidal neurons and specifically target SOM neurons^4^. Long-range connections from cortical^5,6^ or subcortical^7^ areas can activate VIP neurons, which in turn suppress SOM neurons, and disinhibit pyramidal dendrites. This dendritic disinhibitory circuit formed by VIP and SOM neurons is proposed to gate the excitatory inputs targeting pyramidal dendrites^8,9^ (**Fig. 1a**).

**Figure 1.**
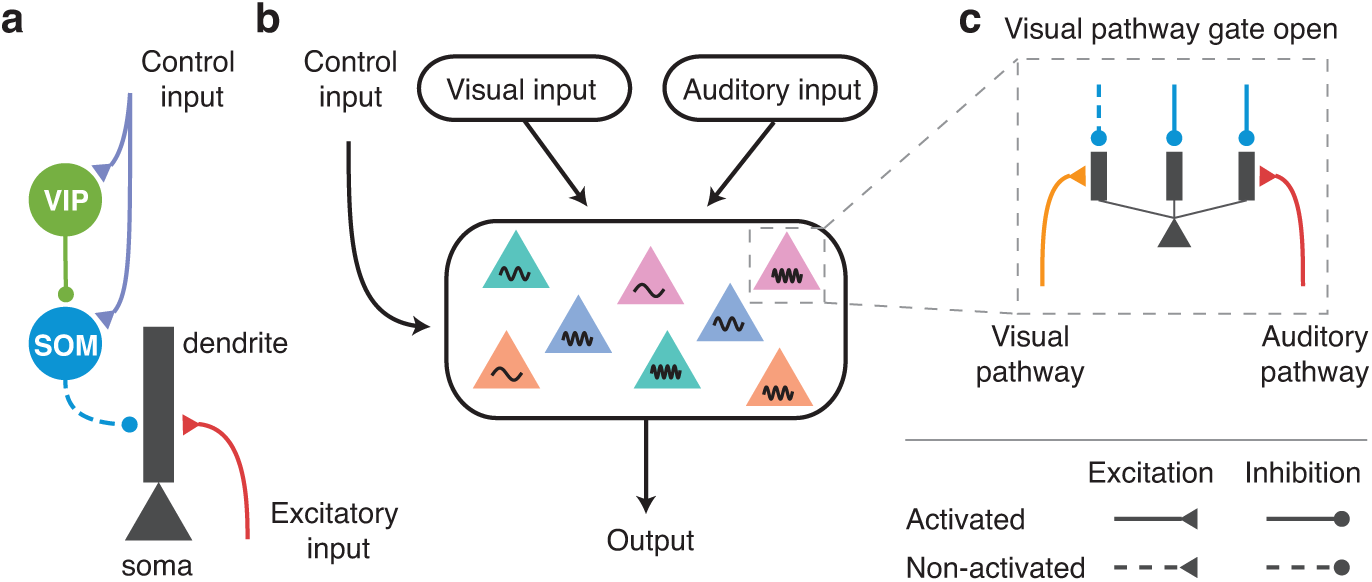
Dendritic disinhibitory circuit as a mechanism for pathway-specific gating. (**a**) Subcellular microcircuit motif for gating through dendritic disinhibition. Dendrites of pyramidal neurons are inhibited by SOM interneurons, which are themselves inhibited by VIP interneurons. A control input (representing a context or a task rule) targeting VIP interneurons (and potentially SOM neurons) can thereby disinhibit pyramidal neuron dendrites, opening the gate for excitatory inputs targeting these dendrites. (**b**) Circuit configuration for pathway-specific gating. Pyramidal neurons receive converging inputs from multiple pathways, e.g. visual and auditory. Single neurons in these areas are selective to multiple stimulus features, indicated here by color and frequency. The processing of each pathway is regulated by the control input. (**c**) Inputs from different pathways target distinct subsets of dendrites of these pyramidal neurons. A pathway can be gated-on by specifically disinhibiting the dendrites that it targets, corresponding to an alignment between excitation and disinhibition. Disinhibition is represented by dashed lines.

Insofar as any cortical area receives inputs from tens of other areas and project to many other areas, information flow across the complex cortical circuit needs to be flexibly gated (or routed) according to behavioral demands. Broadly speaking, there are three types of gating in terms of specificity. First, all inputs into a cortical area may be uniformly modulated up or down. Recent research in mice demonstrated that such gating involves the disinhibitory motif mediated by VIP and SOM interneurons^5,7,10–13^. These studies generally found that VIP neurons are activated, and SOM neurons are inactivated, in response to changes in the animals’ behavioral states, such as when mice receive reinforcement^12^, or start active whisking^5,13^ or running^7^. The reported state change-related activity responses can be remarkably homogeneous across the local population of the same class of interneurons^10,11^.

Second, gating may involve selective information about a particular stimulus attribute or spatial location (for instance, in visual search or selective attention^6^). Whether SOM or VIP neurons are endowed with the required selectivity remains insufficiently known. In sensory cortex, SOM neurons exhibit greater selectivity to stimulus features (such as orientation of a visual stimulus) than PV neurons^14,15^. Furthermore, in motor cortex, SOM neurons have been shown to be highly heterogeneous and remarkably selective for forward versus backward movements (Adler & Gan, *Society for Neuroscience*, 2015).

Third, for a given task, neurons in a cortical area may need to “gate in” inputs from one of the afferent pathways, and “gate out” other afferent pathways^16,17^, which we call “pathway-specific gating”. For instance, imagine yourself sitting in a noisy cafe and trying to focus on your book. Your associational language areas receive converging inputs from both auditory and visual pathways. Opening the gate for the visual pathway while closing the gate for the auditory pathway allows you to focus on reading (**Fig. 1b**). In the classic Stroop task, the subject is shown a colored word, and is asked to either name the color or read the word. One possible solution to this task is for a decision-making area to locally open its gate for the deliberate pathway (color-naming) while closing its gate for the more automatic pathway (word-reading).

Using computational models, we propose that the dendritic disinhibitory circuit can instantiate pathway-specific gating. Each of the many branches of a pyramidal dendrite process its inputs quasi-independently^18^ and nonlinearly^19^. Feedforward and feedback pathways target different regions (e.g. basal or apical tuft) of dendritic trees of pyramidal neurons^20^. We hypothesize that excitatory inputs from different pathways can cluster onto parts of dendrites of pyramidal neurons, which we term “branch-specific” even though inputs from a particular pathway may target multiple branches. This hypothesis is supported by mounting evidence for synap-tic clustering on dendritic branches^21–23^. A pathway can presumably be “gated-on” by specifically disinhibiting the branches targeted by this pathway (**Fig. 1c**), i.e. by a disinhibition pattern aligned with the excitation. This *branch-specific disinhibition* is motivated by findings showing that synaptic inhibition from SOM neurons can act very locally on dendrites, even controlling individual excitatory synapse by targeting the spine^3^ or the pre-synaptic terminal^24^.In this work, we developed a network model with thousands of pyramidal neurons and hundreds of interneu-rons for each (VIP, SOM, and PV) type, and show that pathway-specific gating can be accomplished by the disinhibitory motif, even though the connectivity from SOM neuronstopyramidal neurons is dense: each SOM neuron on average targets more than 60% of neighboring pyramidal neurons (< 200 *μ*m)^25^.

We first characterized how branch-specific disinhibition can efficiently gate excitatory inputs onto pyramidal dendrites. To then test whether the densely-connected interneuronal circuit can indeed support branch-specific disinhibition, we built a dendritic disinhibitory circuit model constrained by experimentally measured single-neuron physiology and circuit connectivity. We found that although SOM-pyramidal connectivity is dense at the level of neurons, at the level of dendrites it is sufficiently sparse to support branch-specific disinhibition, and therefore pathway-specific gating, given that SOM neurons can be selectively controlled. We then showed control inputs targeting both VIP and SOM neurons can selectively suppress SOM neurons as needed. Notably we drew these conclusions under some “worst-case” assumptions to our model such as random interneuronal connectivity. Using a calcium-based synaptic plasticity model, constrained by data, we found that disinhibitory regulation of plasticity can give rise to an appropriate alignment of excitation and disinhibition which is required for pathway-specific gating. Finally, we demonstrated the functionality of this mechanism in a circuit model performing an example context-dependent decision-making task^26^.

Our results suggest that, as an alternative to the proposal that SOM neurons act as a “blanket of inhibition”^27^, they can indeed subserve pathway-specific gating. This work predicts that top-down behavioral control involves rule signals targeting specific interneuron types rather than (or in addition to) pyramidal neurons, and that the disinhibitory motif plays a major role in synaptic plasticity.

## Results

### Pathway-specific gating with dendritic disinhibition

To study dendritic disinhibition, we first built a simplified neuron model with a reduced morphology, constrained to fit physiological data (**Fig. 2a**, **Supplementary Fig. 1**). It comprises one spiking somatic compartment, and multiple dendritic compartments which are electrically coupled to the soma but otherwise independent of each other. The somatic and dendritic compartments have no spatial extent themselves. This choice of morphology is inspired by previous studies showing that different dendritic branches can integrate their local input independently from one another^18^.

**Figure 2.**
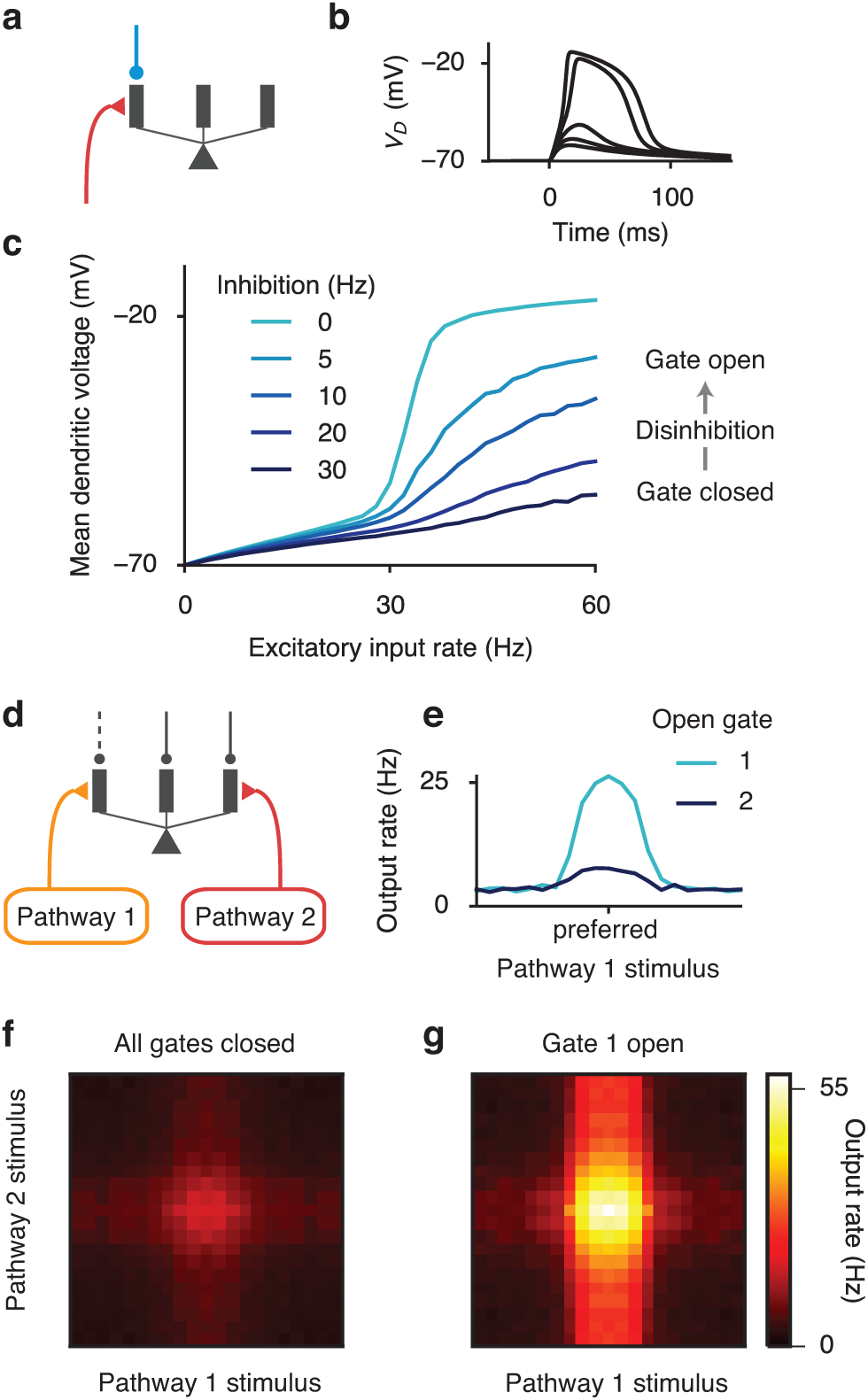
Context-dependent gating of specific pathways. (**a**) A reduced compartmental neuron with a somatic compartment connected to multiple, otherwise independent, dendritic compartments (only three shown). (**b**) Excitatory inputs can generate a local, regenerative NMDA plateau potential in the dendrite. As number of activated synapses increased, there is a sharp nonlinear increase in the evoked dendritic membrane depolarization (*V*_*D*_). (**c**) Disinhibition of the targeted branch opens the gate for the excitatory input. (**d**) A pyramidal neuron receives converging inputs from multiple pathways carrying different stimulus features, giving it selectivity to a preferred stimulus for each feature dimension. Each input pathway targets separate dendrites, which are disinhibited correspondingly in each context by top-down control inputs (not modeled here). (**e**) Tuning curve for input pathway 1, when only this pathway is activated. The input pathway encodes a stimulus feature, e.g. motion direction, with a bell-shaped tuning curve for the input. The preferred feature value corresponds to higher input firing rate. When gate 1 is open by disinhibiting the dendrites targeted by input pathway 1, the neuron exhibits strong tuning (light blue). When gate 2 is instead open, the neuron exhibits weak tuning for the feature (dark blue). The amount of inhibition reduced for a disinhibited dendrite, i.e. the disinhibition level, is 30 Hz. (**f,g**) Two dimensional tuning curves when both pathways are activated. (**f**) In the default context, no dendrites are disinhibited and both pathways are gated off. The neuron exhibits weak responses regardless of the stimulus features. (**g**) When gate 1 is open by disinhibiting branches targeted by pathway 1, the response of this neuron is dominated by tuning to the pathway 1 stimulus, although pathway 2 has a residual impact.

A prominent feature of active processing in thin dendritic branches is their ability to produce NMDA plateau potentials^28^, also called NMDA spikes. The NMDA plateau potential is a regenerative event in which the membrane potential increases nonlinearly and sometimes sharply with the NMDAR input, due to the release of voltage-dependent magnesium block of NMDARs. The reduced neuron model can exhibit NMDA plateau potential in dendrites (**Fig. 2b**), in line with simulations of morphologically reconstructed neuron models (**Supplementary Fig. 1**). The mean dendritic voltage in response to a Poisson spike train input is a sigmoidal function of the input rate, due to the NMDA plateau potential (light blue curve in **Fig. 2c**).

The NMDA plateau potential can be prevented by applying a moderate synaptic inhibition, mediated by GABA_A_ receptors, to the same dendrite (dark blue curve in **Fig. 2c**). Inhibition is particularly effective in controlling this dendritic nonlinearity when excitatory inputs are mediated by NMDA receptors with experimentally-observed saturation, in stark contrast to AMPA receptors (**Supplementary Fig. 1**) or NMDA receptors without saturation (**Supplementary Fig. 2**). Inhibitory input also linearizes the relationship between mean dendritic voltage and excitatory input rate (**Fig. 2c**), due to stochastic transitions into or out of NMDA plateau potential induced by low-rate inhibition (**Supplementary Fig. 2**). Therefore excitatory inputs to a dendritic branch can be efficiently gated by inhibition^29^.

We now consider multiple pathways of inputs targeting distinct sets of dendrites. In the default condition, all dendritic branches receive a high baseline inhibition from dendrite-targeting SOM neurons^5,13^, closing gates for all pathways. Disinhibiting the branches targeted by one pathway can selectively open the gate for this pathway while keeping the gates closed for other pathways (**Fig. 2d**). When a gate is open, the neuron’s output firing rate transmits the stimulus selectivity of the corresponding input pathway most effectively (**Fig. 2e**).

When two excitatory pathways are activated simultaneously, we can plot the neuron’s response to stimulus variables of both pathways, i.e. the two-dimensional tuning curve (**Fig. 2f,g**). In the default condition when all gates are closed, there is little response to either pathway (**Fig. 2f**). By specifically disinhibiting the branches targeted by pathway 1, we can open the gate for pathway 1. With gate 1 opened, the neuron is primarily selective to pathway 1 stimuli (**Fig. 2g**). The remaining impact of pathway 2 stimuli is due to the fact that the impact of excitatory inputs can never be fully counteracted by dendritic inhibition.

The gating mechanism worsens when a fraction of excitatory input is mediated by AMPARs, but improves when a fractionofinhibitoryinputismediatedby GABA_B_ receptors (**Supplementary Fig. 3**). Under *in vivo* conditions, the relative contribution of AMPAR-mediated inputs is likely quite low, as a result of a lower glutamate affinity and a stronger desensitization^19^. For parsimony, in the following sections, excitatory synaptic inputs are mediated only by NMDARs which are critical to the nonlinear dendritic computations, and inhibitory inputs are mediated only by GABA_A_Rs.

### Performance of gating in pyramidal neurons

Which circuit properties determine the effectiveness of pathway-specific gating in our model? A neuron responds to its optimal stimulus from an input pathway with (baseline-corrected) firing rate (*r*_on_) when the pathway is gated-on, and (*r*_off_) when the pathway is gated-off, which could be readily measured experimentally. The gating selectivity is then quantified,

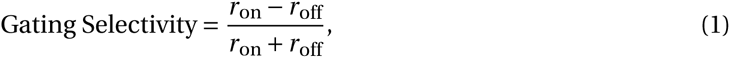

which ranges from 0 (no gating) to 1 (perfect gating). We developed a multi-compartmental rate model^18^ that greatly improves the efficiency of the circuit model simulation. The rate model is fitted to quantitatively reproduce the activity of the spiking neuron model (**Supplementary Fig. 4**, see Supplemental Information for details).

We first tested how gating selectivity depends on our assumption of branch-specific disinhibition in a single-neuron setting. Here we assume an alignment of excitation and disinhibition patterns, which can be achieved through synaptic plasticity as shown later. Each excitatory pathway targets *N*_*disinh*_ randomly chosen dendrites, out of *N*_*dend*_ total dendrites, and this pathway is gated-on by specifically disinhibiting these same *N*_*disinh*_ dendrites (**Fig. 3a**). Due to the random independent selection of targeted dendrites for each pathway, inputs from two different pathways often overlap.

**Figure 3.**
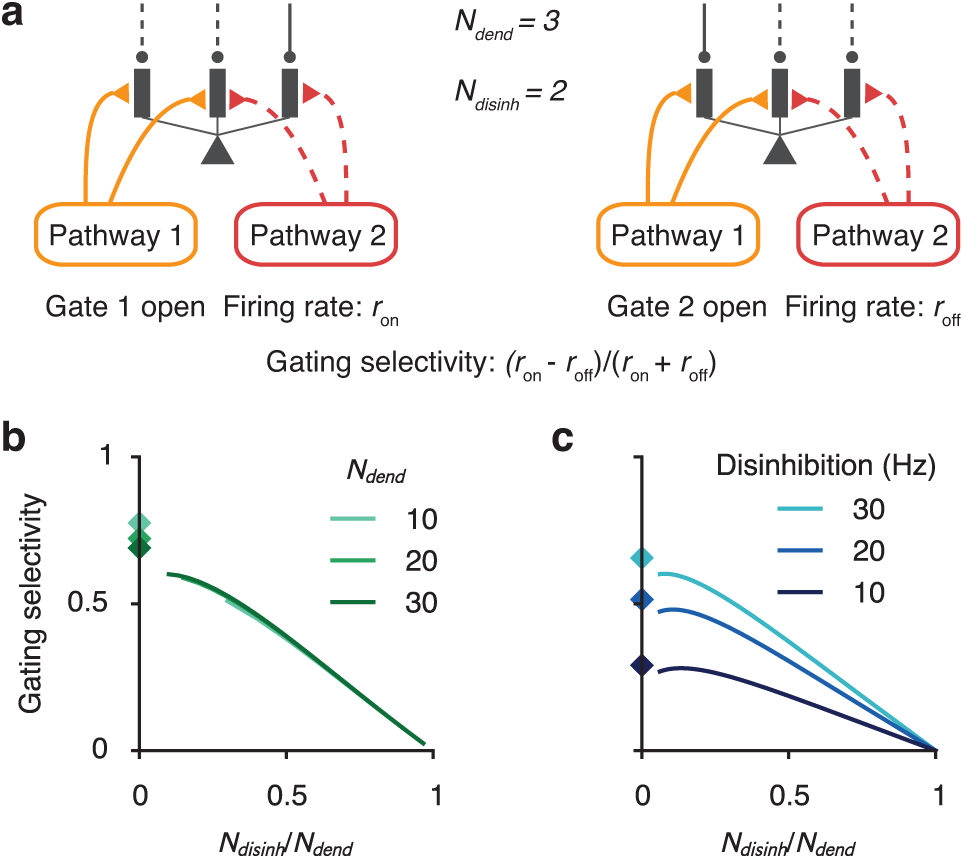
Characterization of gating selectivity in pyramidal neurons. (**a**) Schematic of gating, presenting pathway 1 input when gate 1 is opened (left) or gate 2 is opened (right). There are *N*_dend_ available dendrites in total. Each input pathway targets *N*_disinh_ dendrites. To gate a pathway on, these exact *N*_disinh_ dendrites are disinhibited, creating an aligned pattern of disinhibition. Each pathway selects dendrites randomly and independently from other pathways, which can result in overlap of the excitation-disinhibition patterns across pathways. When *N*_disinh_ is large, projections from different pathways are more likely to overlap. The neuron’s firing rate is *r*_on_ and *r*_off_ in response to the preferred stimulus of the gated-on (left) and gated-off (right) pathway respectively. The gating selectivity is defined as (*r*_on_ − *r*_off_)/(*r*_on_ + *r*_off_), which is 1 for perfect gating and 0 for no gating. (**c**) Gating selectivity increases as excitation/disinhibition patterns become sparser, i.e. with a smaller proportion of targeted and disinhibited dendrites for a pathway (*N*_disinh_/*N*_dend_). Diamonds mark the case of non-overlapping excitatory projections, corresponding to the limit of maximal sparseness. (**d**) Gating selectivity is higher with stronger disinhibition, for all sparseness levels.

We found that gating selectivity depends critically on the sparseness of the disinhibition (**Fig. 3b**), defined as the proportion of targeted/disinhibited dendrites *N*_*disinh*_/*N*_*dend*_. Gating selectivity improves when disinhibition patterns are sparsened, because the proportion of dendrites that receive overlapping inputs is reduced. We can approximate the limit of *N*_*disinh*_/*N*_*dend*_ → 0 with non-overlapping disinhibition pattern (diamonds in **Fig. 3b,c**). In this case, the gating selectivity is highest but below 1, due to the remaining impact of inputs targeting inhibited dendrites, and is therefore modulated by the level of disinhibition (**Fig. 3c**).

### Pathway specific gating in an interneuronal circuit of SOM neurons

We have shown that a key determinant of gating performance is the sparseness of innervation patterns onto the dendritic tree. Yet the connectivity from SOM interneurons to pyramidal neurons is dense^25^. Is it possible for the proposed gating mechanism to function in a cortical micro-circuit constrained by the dense interneuronal connectivity? To address this issue, we built an interneuronal circuit model, containing hundreds of VIP and SOM interneurons and thousands of pyramidal neurons, constrained by anatomical and physiological data. We considered “worst-case” conditions in which interneuronal connectivity is completely random (as our gating mechanism can be facilitated by structured connectivity). Surprisingly, we found that relatively high gating performance is achievable under these conditions. We analyzed gating in this circuit in two steps: First, assuming SOM neurons are context-selective, we characterized how the SOM-pyramidal sub-circuit can support high gating selectivity. Second, we characterized how SOM neurons can become context-selective in the VIP-SOM-pyramidal circuit.

First, we built a simplified model of a SOM-pyramidal sub-circuit (**Fig. 4a**), which corresponds roughly to a cortical L2/3 column (400*μm* × 400/*μm*). The model contains *N*_pyr_ (≈ 3,000) multi-compartmental pyramidal neurons, each with *N*_dend_ (≈ 30) dendrites, and *N*_SOM_ (≈ 160) SOM neurons (see **Supplementary Table 1**). Here we analyze the dependence of gating selectivity on the connectivity from SOM to pyramidal neurons. We consider worst-case conditions in which these connections are random, subject to the SOM-to-pyramidal connection probability of *P*_SOM→pyr_ (≈ 0.6). Assuming that a SOM neuron chooses to target each pyramidal dendrite independently with a SOM-to-dendrite connection probability of *P*_SOM→dend_, then we have

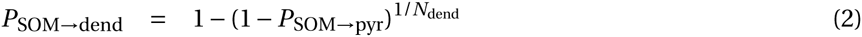

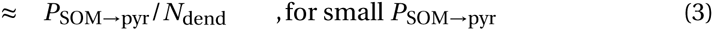

**Figure 4.**
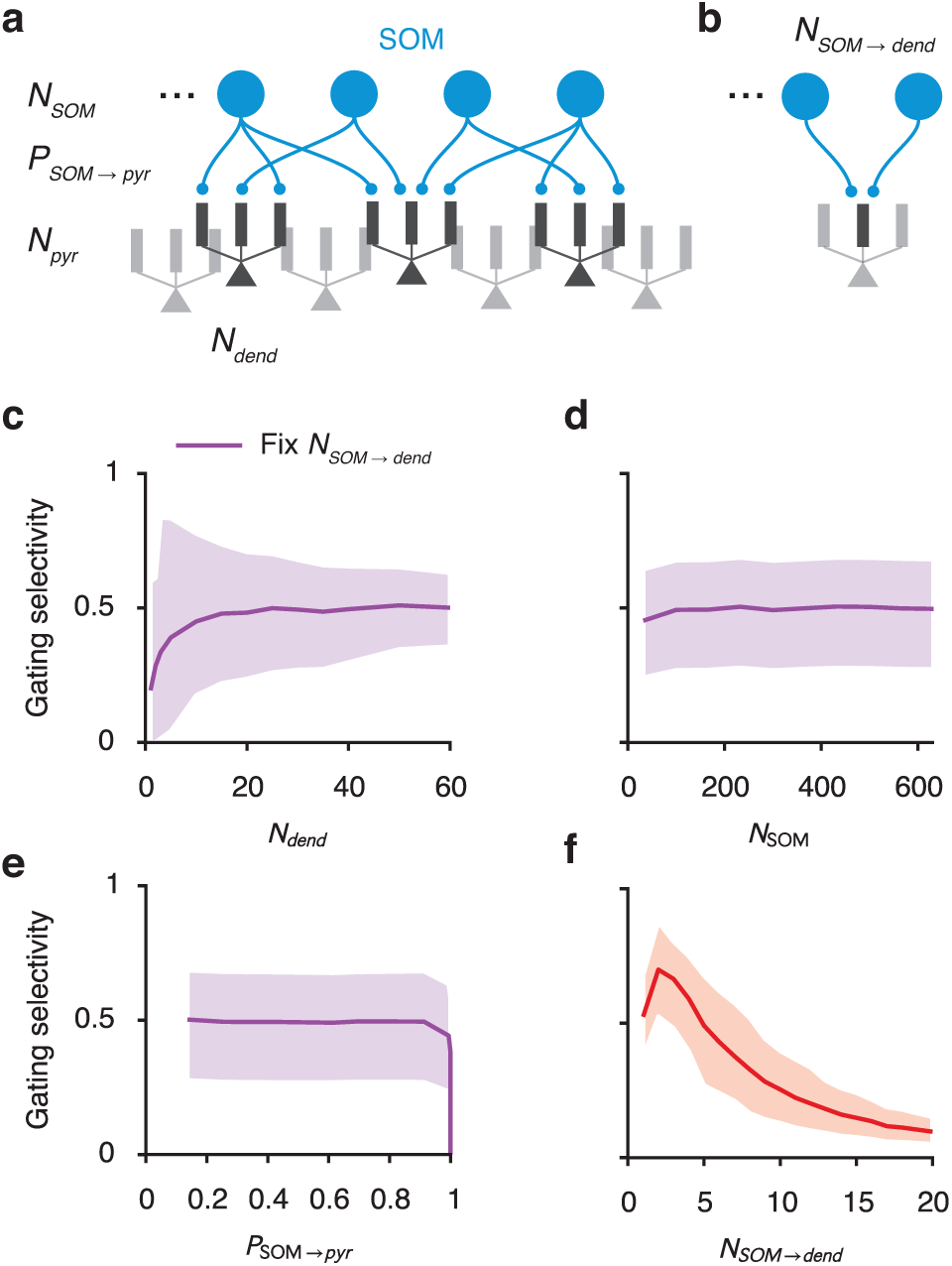
Gating selectivity as functions of SOM-pyramidal circuit parameters. (**a**) A biologically-constrained model for a cortical column of SOM and pyramidal neurons. We only modeled the SOM-to-pyramidal connections. The model is subject to experimentally-measured constraints of the following parameters: number of SOM neurons (*N*_SOM_), connection probability from SOM to pyramidal neurons (*P*_SOM→pyr_), and the number of dendrites on each pyramidal neuron (*N*_dend_). We consider the “worst case” scenario that the SOM-to-dendrite connections are random. Finally we assume for now that control input for each pathway suppresses a random subset of SOM neurons. The different contrasts used are for illustration purpose only. (**b**) A critical parameter for the SOM-to-pyramidal circuit is the number of SOM neurons targeting each dendrite (*N* _SOM→dend_). This parameter can be calculated using other experimentally-measured parameters under the assumption of random connectivity, *N* _SOM→dend_ = *N*_SOM_ · [1 − (1 -*P*_SOM→pyr_)^1/*N*_dend_^]. (**c-e**) Gating selectivity only weakly depends on *N*_dend_ (**c**), *N*_SOM_ (**d**), and *P*_SOM→pyr_ (**e**) if *N*_SOM→dend_ is kept constant by co-varying another parameter. The plotted curve marks the mean and the shaded region marks the bottom 10% to top 10% of the neuronal population. (**f**) Gating selectivity is high when each dendrite is targeted by a few SOM neurons. Given experimental measurements of *P*_SOM→pyr_ ≈ 0.6,*N*_dend_ ≈ 30,*N*_SOM_ ≈ 160, we obtained *N*_SOM→dend_ ≈ 5, leading to relatively high gating selectivity ~ 0.5. Total strength of inhibition onto each pyramidal dendrite is always kept constant when varying parameters.

Under this assumption, a SOM neuron on average targets *N*_dend_·*P*_SOM→dend_/*P*_SOM→pyr_ ≈ 1.5 den-drites of a pyramidal neuron given that the two are connected. Each SOM-dendrite connection can correspond to multiple (3-5) clustered synapses^30^. So each SOM neuron can make on average 5-8 synapses onto a pyramidal neuron. The connection probability between two neurons is higher at closer proximity^25^, leading to a even higher number of contacts.

In a default state, SOM neurons fire at a relatively high baseline rate around 10 Hz^5,13^, closing the gates to all inputs. To open the gate for pathway 1, a randomly chosen subset (50%) of SOM neurons are suppressed, resulting in a pattern of disinhibition across dendrites. Again we assume the excitatory input pattern of pathway 1 is aligned with the corresponding disinhibition pattern. Notably, disinhibition patterns for different pathways generally overlap due to the random selection of SOM neurons and the random connectivity. This overlap can be reduced with either structured connections or inhibitory plasticity.

Under the above assumptions, the circuit achieves a mean gating selectivity around 0.5, equivalent to *r*_on_ ≈ 3*r*_off_. We found that the impact of these circuit parameters is determined by one critical parameter: the number of SOM neurons targeting each dendrite *N*_SOM→dend_ = *N*_SOM_ · *P*_SOM→dend_ ≈ 5 (**Fig. 4b**, see also Supplemental Mathematical Appendix). When we vary parameters while keeping *N*_SOM→dend_ fixed, the gating selectivity remains largely constant (**Fig. 4c-e**). We found that gating selectivity is highest when *N*_SOM→dend_ is small (**Fig. 4f**), and decreases as we increases *N*_SOM→dend_. Because the overall strength of inhibition has a simple effect on the gating selectivity (**Fig. 3c**), we keep it fixed when varying other parameters.

Each dendrite should more appropriately be interpreted as an independent computational unit. When inhibitory connections control individual excitatory connection through pre-synaptic receptors^24^ or by targeting spines^3^, the independent unit would be single excitatory synapses. This leads to a lower effective value of *N*_SOM→dend_, then a higher gating selectivity.

### Pathway specific gating in an interneuronal circuit of SOM and VIP neurons

Having analyzed the SOM-pyramidal connectivity, we next examined how SOM neurons can be context-selective, and characterized the gating selectivityin acircuit model containing VIP, SOM, and pyramidal neurons. On top of the previous SOM-pyramidal sub-circuit, We added *N*_VIP_ VIP neurons that only target SOM neurons^4^. Here we assume VIP neurons target all SOM neurons with connection probability *P*_VIP→SOM_. Broadly speaking, we found two scenarios in which SOM neurons can be suppressed selectively based on the context, depending on the targets of the top-down or locally-generated control inputs (**Fig. 5**).

**Figure 5.**
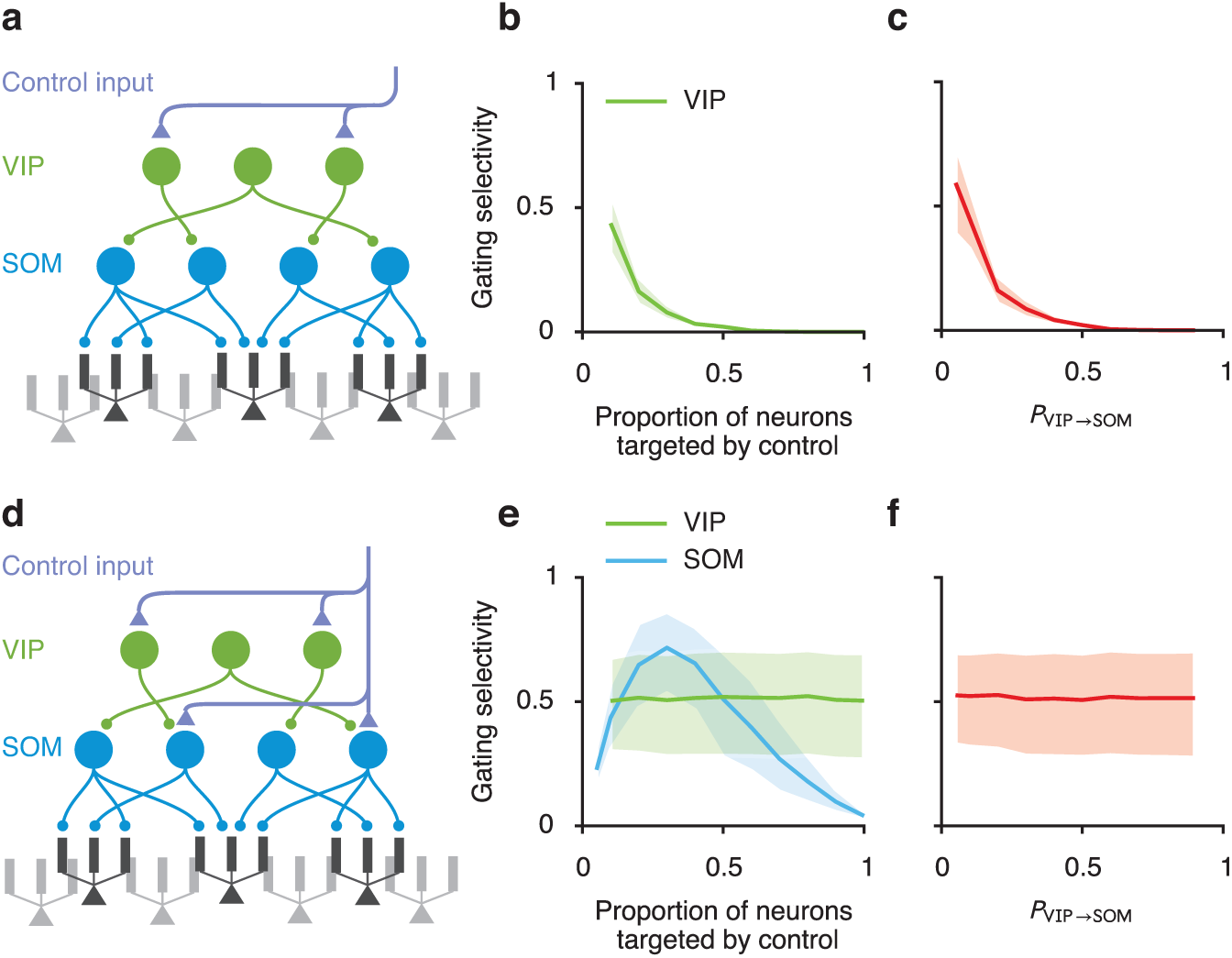
Two possible schemes of control for the interneuronal circuit. We built a simplified circuit model containing VIP, SOM, and pyramidal neurons. (**a-c**) Control signals target only VIP neurons. (**a**) In this scheme, for each pathway, control inputs target a random subset of VIP neurons. And the connection probability from VIP to SOM neurons is *P*_VIP→SOM_. (**b,c**) Good gating selectivity is only achieved when a small subset of VIP neurons is targeted by control inputs (**b**), and when the VIP-SOM connections is sparse (**c**). (**d-f**) Control signals target both VIP and SOM neurons. (**d**) In this scheme, we assume that for each pathway control inputs target a random subset of VIP and SOM neurons. (**e**) Gating selectivity depends on the proportion of SOM (blue) but not VIP (green) neurons targeted by control input. (**f**) Gating selectivity does not depend on *P*_VIP→SOM_. Curves and shaded regions are as in **Fig. 4**.

In the first scenario, control inputs target VIP neurons solely (**Fig. 5a**). In this intuitive scenario, control inputs excite VIP neurons, which in turn inhibit SOM neurons thereby disinhibiting pyramidal dendrites. Gating selectivity is high only if a small proportion of VIP neurons is targeted by control (**Fig. 5b**), indicating that VIP neurons must be context-selective, and VIP-SOM connections need to be sparse (**Fig. 5c**). VIP-SOM connectivity could possibly be effectively sparse on the scale of a cortical column, since the axonal arbor of VIP neurons are rather spatially restricted^31^. When varying parameters, we kept fixed the overall baseline inhibition received by each SOM neuron and the overall strength of control inputs.

In the second scenario, excitatory control inputs target both VIP and SOM neurons (**Fig. 5d**). If the VIP-SOM connectivity is dense, then VIP neurons activated by control inputs will provide nearly uniform inhibition across all SOM neurons (**Supplementary Fig. 5**). However, SOM neurons can receive selective excitation if the control inputs only directly target a subset of SOM neurons. If the inhibition is on average stronger, then the overall effect is a selective suppression of SOM neurons (**Supplementary Fig. 5**). As a result, gating selectivity no longer depends on the proportion of VIP neurons targeted by control inputs, but does depend on the proportion of SOM neurons targeted (**Fig. 5e**). Therefore SOM neurons are context-selective, but VIP neurons need not be. Similarly, gating selectivity does not depend on the connection probability from VIP to SOM neurons, *P*_VIP→SOM_ (**Fig. 5f**).

In summary, in order to achieve branch-specific disinhibition, control inputs targeting interneurons have to be selective. Notably, the level of specificity required for the control inputs depends strongly on the neurons they target. When targeting only VIP neurons, the control inputs have to be highly selective (**Fig. 5a,b**). However, when control inputs target both VIP and SOM neurons, high gating selectivity can be achieved in a much broader range of parameters, reducing the level of specificity required (**Fig. 5d,e**).

### Pathway specific gating in an interneuronal circuit of SOM, VIP, and PV neurons

PV neurons receive inhibition from themselves and SOM neurons, and project to perisomatic areas of pyramidal neurons^1^. Suppression of SOM neurons therefore also leads to disinhibition of PV neurons and an increase of somatic inhibition onto pyramidal neurons. We included PV neurons into our interneuronal circuit model (**Fig. 6a**), and found that this inclusion and the consequent increase in somatic inhibition strictly improve gating selectivity in a wide range of parameters (**Fig. 6b**). Since the SOM-to-PV and PV-to-pyramidal neuron connections are dense^27^, a selective pattern of SOM suppression will result in an elevated somatic inhibition that is almost uniform across pyramidal neurons (**Supplementary Fig. 6**). Furthermore, we proved that an uniform increase in somatic inhibition will always improve gating selectivity, except when the somatic inhibition is unreasonably strong (see **Supplemental Mathematical Appendix**).

**Figure 6.**
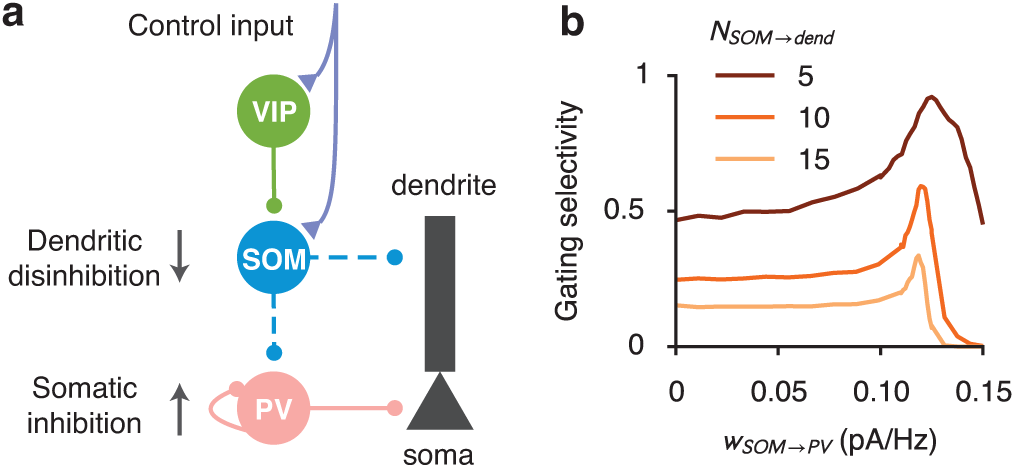
Somatic inhibition improves gating selectivity. (**a**) PV neurons project to the somatic areas of pyramidal neurons, and are inhibited by SOM neurons and themselves. Suppression of SOM neurons cause disinhibition of PV neurons, therefore an increase in somatic inhibition onto pyramidal neurons. (**b**) A moderate increase in somatic inhibition always improves gating selectivity. We included PV neurons and their corresponding connections in the model of **Fig. 5d**. Gating selectivity increases as a function of the SOM-to-PV connection weights (*w*_SOM→PV_) in a wide range (see Supplementary Mathematical Appendix for a proof). However, when gating selectivity is low without PV neurons (light curve), the peak of this increase is lower and the slope is sharper. Gating selectivity starts to decrease when the SOM-to-PV connection, therefore the somatic inhibition, is too strong that the responses of many pyramidal neurons are completely suppressed.

For an intuitive explanation, consider a linear input-output function in the soma. Gating selectivity is based on the relative difference between the pyramidal neuron responses when the gate is open (*r*_on_) and when the gate is closed (*r*_off_). Providing an equal amount of somatic inhibition in these two conditions is equivalent to subtracting both values by the same constant, which will enhance the relative difference.

### Inhibitory modulation of synaptic plasticity and learning pathway-specific gating

A critical feature of our scheme is the alignment between excitation and disinhibition patterns (**Fig. 1c**): pyramidal dendrites targetedbyanexcitatoryinput pathway are also disinhibited when the gate is open for that pathway. Dendritic disinhibition can regulate synaptic plasticity^32,33^. We hypothesized that such an alignment can naturally arise as a result of the regulated plasticity. To test this hypothesis, we first established a realistic calcium-based plasticity model for dendrites in our reduced spiking neuron model. Pre-and post-synaptic spikes induce calcium transients in dendrites, which determine the synaptic weight changes^34^ (**Fig. 7a**). We fitted parameters of the model to capture experimental data^35^ (**Supplementary Fig. 7**). Our model also quantitatively predicts findings that were not used in the fitting.

**Figure 7.**
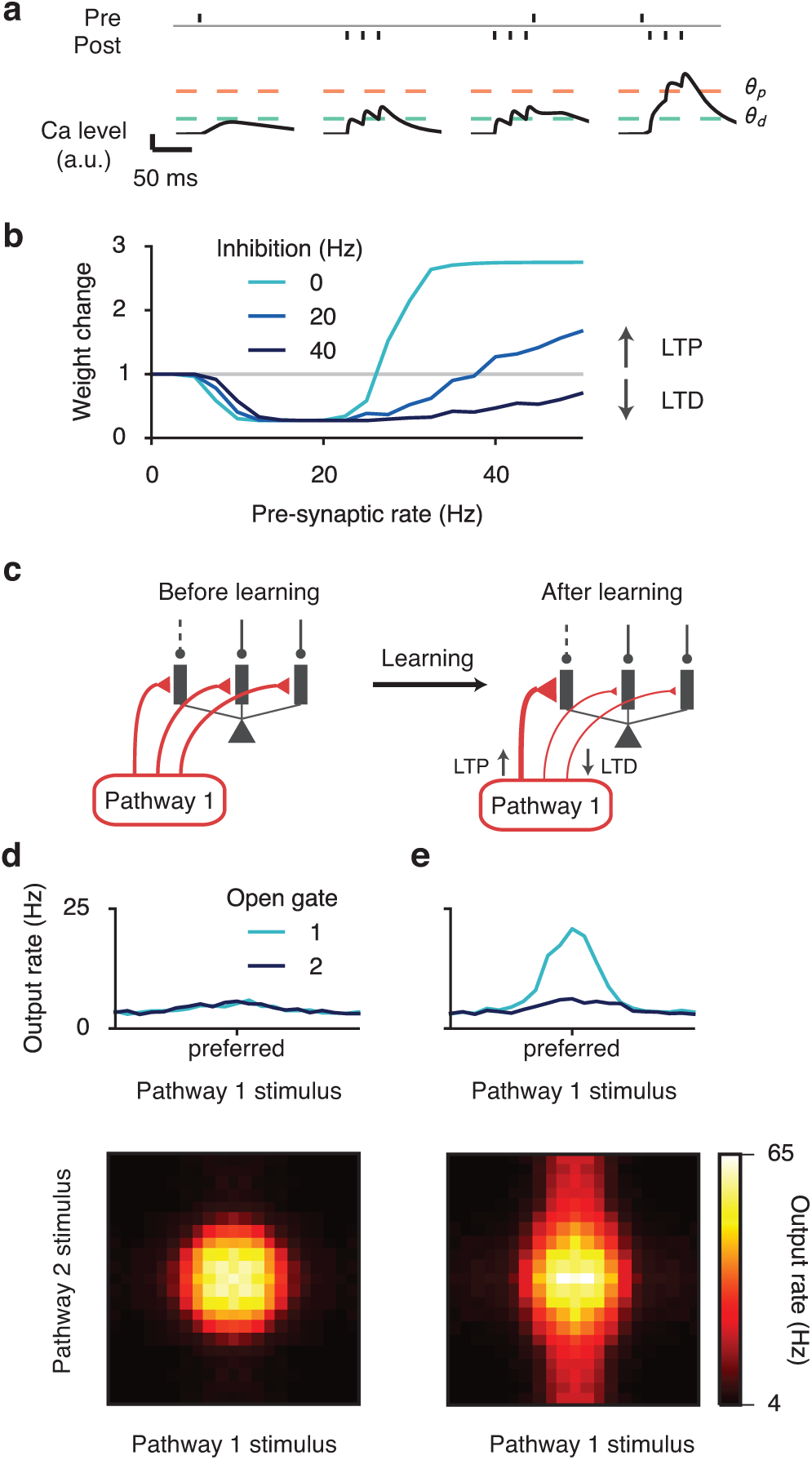
Learning to gate specific pathways. (**a**) Model schematic. Pre-and post-synaptic spikes both induce calcium influx. The overall synaptic weight change is determined by the amount of time the calcium level spends above thresholds for depression (*θ*_*d*_) and potentiation (*θ*_*p*_)^34^. The model is fitted to experimental data, and is able to quantitatively predict results not used in the fitting (**Supplementary Fig. 7**). (**b**) Dendritic inhibition makes potentiation harder to induce. With background-level inhibition (light blue), synaptic weight change shows three regimes as a function of excitatory input rate: no change for low rate, depression for medium rate, and potentiation for high rate. With a medium level of inhibition (dark blue), potentiation requires a higher excitatory input rate. With relatively strong inhibition (black), potentiation becomes impossible within a reasonable range of excitatory input rates. The post-synaptic rate is fixed at 10 Hz. (**c**) Learning paradigm. (Left) Excitatory synapses from each pathway are initialized uniformly across dendrites. When pathway 1 is activated, specific branches of the neuron are disinhibited (dashed line), i.e. gate 1 is open. During learning, only one pathway is activated at a time. (Right) After learning, activated excitatory synapses onto the disinhibited branches are strengthened, while activated synapses onto inhibited branches are weakened, resulting in an alignment of excitation and disinhibition patterns. Synaptic weights of non-activated synapses remain unchanged (not shown). (**d**) Response properties of the neuron before learning. (Top) Tuning curve of the neuron when only pathway 1 is presented. The neuron shows no preference to the gate opened prior to learning. (Bottom) Two-dimensional tuning curve of the neuron when both pathways are simultaneously presented and gate 1 is open. See **Fig. 2** for the definition of the tuning curves. (**e**) Response properties of the neuron after learning. (Top) The neuron shows strong tuning to pathway 1 input when gate 1 is open. (Bottom) When both pathways are presented, the neuron’s response is primarily driven by pathway 1 stimulus, although pathway 2 stimulus also affects the neuron’s firing.

The calcium-based plasticity model allows us to naturally study the effects of dendritic disinhibition on synaptic plasticity and their functional implications. Again we assume that pre-and post-synaptic firings are Poisson spike trains with specified rates. We found that dendritic inhibition can shift the plasticity from potentiation to depression, even when the pre-synaptic excitatory input rate and the post-synaptic firing rate are both kept constant (**Fig. 7b**), consistent with previous modeling findings^33^. We note that plasticity models based solely on pre-and post-synaptic neuronal firing would not predict the inhibitory modulation of synaptic plasticity.

We then tested whether disinhibitory regulation of plasticity can support the development of excitation-disinhibition alignment, as needed for pathway-specific gating (**Fig. 7c**). Importantly, the strength of disinhibition is realistic, similar to those used throughout this paper. Initially, excitatory synapses from each pathway are uniformly distributed across the dendritic branches of single neurons. Different excitatory pathways are then activated one at a time. Whenever a pathway is presented, a particular subset of dendrites are disinhibited, while the rest of the dendrites remain inhibited. Through calcium-based excitatory plasticity, the activated excitatory synapses targeting the disinhibited dendrites become strengthened, whereas those targeting the inhibited dendrites become weakened. Synapses not activated remain the same regardless of the inhibition level (see **Fig. 7b**). After learning, the alignment of excitation and disinhibition patterns support pathway-specific gating (**Fig. 7d,e**; compare with **Fig. 2e**), with a gating selectivity around 0.7. These findings show that a key aspect of the gating architecture, namely the alignment of excitation and disinhibition patterns, can emerge naturally from the interaction between excitatory synaptic plasticity and context-dependent disinhibition.

### Modeling a flexible behavior with pathway-specific gating

How is gating at the neural level related to gating at the behavioral level? Is moderate gating selectivity (e.g., ~0.5 as above) sufficient to explain performances in flexible cognitive tasks? To address these issues, we applied our model to a context-dependent decision-making task^26^. In this task, the behavioral response should be based on either the motion direction or the color of a random-dots motion stimulus, depending on the context cued by a rule signal (**Fig. 8a**).

**Figure 8.**
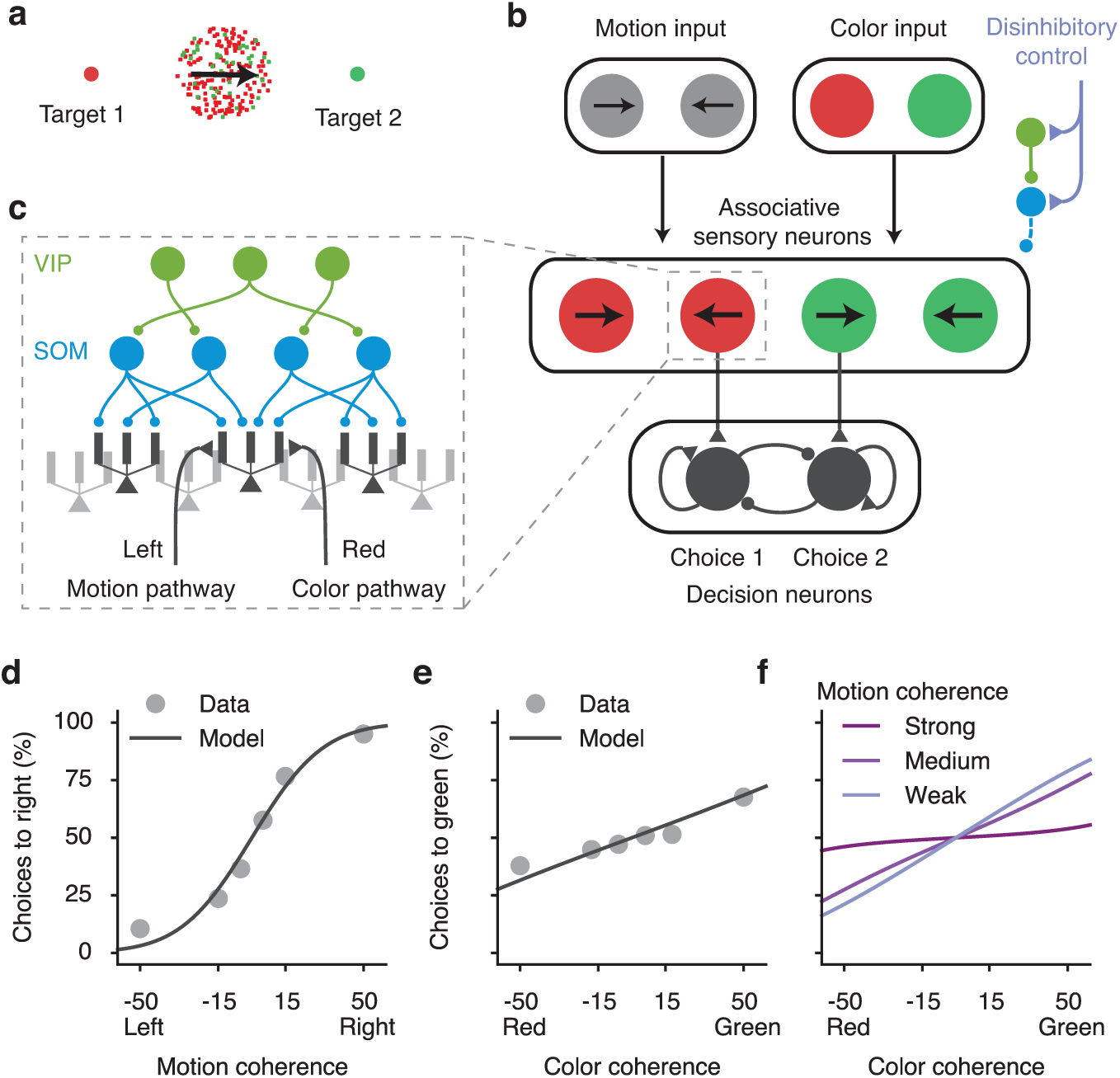
Pathway-specific gating in an example context-dependent decision-making task. (**a**) A flexible decision-making task. Depending on the context, subject’s behavioral response should be based on either the color or the motion direction of the stimulus. (**b**) The circuit model scheme. Motion and color pathways target associative-sensory neurons, which are subject to context-dependent disinhibitory control. Neurons preferring color and motion evidence for the same target project to the corresponding neural pool in the decision-making circuit. (**c**) Associative-sensory neurons receive converging inputs from both motion and color pathways, and are controlled by the dendrite-targeting interneuronal circuit. (**d-f**) Fit and prediction of behavioral performance. Behavioral performance in the motion context as a function of motion coherence (**d**) and color coherence (**e**) for a monkey (dots), and the model’s fit (line). Experimental data are extracted from [26]. The model can capture the behavioral performance of a monkey. (**f**) In the model, impact of the irrelevant pathway (color) is strongest when the relevant pathway signal is weak (with low motion coherence).

We built a stylized neural circuit model to implement this task using pathway-specific gating through dendritic disinhibition (**Fig. 8b**). The local circuit comprises a sensory network and a decision network. The sensory network contains pyramidal neurons that receive convergent sensory inputs from both motion and color pathways, and they group into four pools according to their selectivities to color and motion evidence. The dendrites of pyramidal neurons are controlled by the VIP-SOM interneuronal circuit described above (**Fig. 8c**, **5d**). A subset of pyramidal neurons with high gating selectivity projects to the decision network. Pyramidal neurons representing color and motion evidence for the same target project to the corresponding decision neural pool. The decision network, as modeled previously^36^, is a strongly recurrent network that generates a winner-take-all decision based on its inputs.

We fitted the performance of the model to a monkey’s psychometric behavioral data from [26], using three free parameters in the model, namely the proportion of sensory neurons that project to the decision network, and the overall connection strengths from the input pathways to the sensory network and from the sensory network to the decision network. By fitting these three parameters, we obtained a quantitative match of the empirical psychometric performance, as a functionofrelevant (**Fig. 8e**) and irrelevant (**Fig. 8f**) features. Our model shows that the impact of the irrelevant information should be stronger when the relevant information is more ambiguous (with lower motion coherence, for instance) (**Fig. 8g**). Although at its default parameters the interneuronal circuit model can show similar task performance as the empirical data, we found that it can no longer fit the empirical performance if we significantly degrade the neural gating selectivity (**Supplementary Fig. 8**). This simulation therefore serves as a proof-of-principle to demonstrate the potential of dendritic disinhibition as a mechanism for pathway-gating, and as a link to assess the utility of neural gating selectivity in terms of flexible behavioral performance.

## Discussion

A canonical cortical microcircuit motif specialized for disinhibition of pyramidal neuron den-drites was proposed theoretically^8^ and has received strong empirical support from a series of recent experiments^4–7,12,13,37^. Here we explored the functional roles of dendritic disinhibition using computational modeling, at both the single-neuron and circuit levels. In contrast to somatic disinhibition, dendritic disinhibition can gate the inputs to a neuron^8,29^. We propose that dendritic disinhibition can be utilized to gate inputs from separate pathways, by specifically dis-inhibiting dendrites that receive inputs from a target pathway.

We studied the effectiveness of gating in an interneuronal circuit constrained by experimental data, which have become available only in recent years thanks to the advance of optogenetics and other experimental tools. Where data are not available, we considered the “worst-case scenario”, namely, connections from VIP to SOM neurons, and from SOM to pyramidal dendrites are completely random, which is most likely not the case^38^ and any specificity would facilitate our proposed mechanism. Although the SOM-to-pyramidal connections are dense, we found that the connectivity from SOM neurons to pyramidal dendrites are actually sparse enough to support branch-specific disinhibition. We found that the increase of somatic inhibition mediated by the SOM-PV-pyramidal neuron connections further improves gating selectivity. We demonstrated that branch-specific clustering of excitatory pathways can naturally emerge from disin-hibitory regulation of synaptic plasticity. As proof of principle, we applied this mechanism to a model for a recent experiment using a context-dependent decision-making task^26^.

Inhibitory connections in cortex tend to be dense^25^. This finding has led to the proposal that cortical inhibition functions as a locally non-selective “blanket of inhibition”^27^. In contrast, our study offers an alternative perspective, which is compatible with dense interneuronal connectivity and has very different implications for circuit functions. The dense connectivity is measured on a cell-to-cell level. Nonetheless, connections from dendrite-targeting SOM interneurons can be sparse at the level of the dendritic branch, and therefore potentially selective as required for our gating scheme. Our alternative proposal is fundamentally grounded in consideration of dendritic branches as functional units of computation^18^.

### Circuit requirements for pathway-specific gating

A key assumption and prediction of our model is clustering of excitatory pathways onto pyramidal neuron dendritic branches. The computational benefits of input clustering have been previously proposed^39^. There is mounting experimental evidence for input clustering, from anatomical and physiological studies^21,22^[for a review see 23]. Consistent with our model, experimental studies have shown that input clustering can emerge through NMDAR-dependent synaptic plasticity^40^, and that clustering is functionally related to learning^22,41^. Our model assumption that branch-specific clustering occurs at the level of pathways remains to be directly tested.

Another feature necessary for our mechanism is the alignment between excitation and disinhibition, which we found can be achieved through synaptic plasticity on excitatory synapses. This feature could also potentially be achieved through inhibitory plasticity^42^, by adapting the disinhibition pattern to align with fixed excitatory inputs. These two forms of plasticity are complementary, and both are likely at play. Indeed, a recent study found that during motor learning, spine reorganization on dendrites of pyramidal neurons is accompanied by change in the number of SOM-neuron synapses onto these dendrites^38^. One appeal of studying excitatory plasticity here is that our calcium-based plasticity model could be quantitatively constrained by data and therefore tested in a biologically plausible regime. At present, much lessisknown experimentally about the dependence of inhibitory plasticity on pre-and post-synaptic spike timing, dendritic calcium levels, or the class of interneuron^43^.

### Model predictions

Our model makes specific, experimentally testable predictions. One of the most straightforward and testable predictions is that SOM neurons should show context/rule selectivity in some context-dependent or rule-based tasks. Surprisingly, we found that VIP neurons need not be context-selective, as long as SOM neurons are directly receiving context-selective excitatory control inputs (**Fig. 5d-f**). Experimental disruption of these context-selective interneurons should impair the animal’s ability to perform context-or rule-dependent choice tasks. The context-selectivity of SOM or VIP neurons is not necessarily presented in every behavioral task. For instance, a recent study, recording in mouse prefrontal cortex during a auditory discrimination task, found highly homogeneous responses within SOM and VIP populations^10^. We propose that SOM neurons are more likely to exhibit selectivity to context or task in experiments in which the animal performs multiple tasks and branch-specific dendritic spikes also exhibit task selectivity [e.g.,44, Adler & Gan, *Society for Neuroscience*, 2015]. A direct test of our model awaits future experiments in a task-switching paradigm in order to examine gating of different pathways into association cortical areas and the selective changes of activity in SOM neural subpopulations. We emphasize that interneuron classes in our model should be more appropriately interpreted according to their projection targets rather than their biochemical markers.

Branch-specific dendritic spikes are already observed experimentally, and SOM neurons are critical for this branch-specificity^44^. It is however unknown whether SOM-mediated inhibition is also branch-specific. Direct patch-clamping of pyramidal-neuron dendrites *in vivo*^45^ can isolate inhibitory currents on individual branches, and provide a direct test for our hypothesis, although such an experiment is technically difficult at present. Our plasticity model predicts that SOM interneurons play a critical role in the learning-related emergence of branch-specific clustering of excitatory synapses on pyramidal neuron dendrites^22^.

### Relation to other gating models

Flexible gating, or routing, of information has been a long-standing problem in computational neuroscience^46^, for which a number of models have been proposed. Among proposed ideas are dynamic synaptic weight modulation^47^, gain modulation^46^, synchrony in the input signals^16^, perfect balance of excitation and inhibition^17^, up/down state-switch in dendrites^48^, switching between different neural pools that receive inputs from distinct pathways^49^, and rule signaling as a selection vector^26^. Gating cortico-cortical communication has also been proposed as a role of the basal ganglia^50^. Notably, most of these models implement a form of soft gating, which modulates the effective strength of incoming pathways instead of performing a binary on-off switch on them.

These prior models did not exploit the computational power of dendrites (except for [48]) or the roles of specialized classes of interneurons. Harnessing dendrites rather than populations of intermediate neurons saves the number of neurons needed by many-fold. Only in the limit of one dendrite per pyramidal neuron does our mechanism become conceptually similar to gating mechanisms operating on the neuronal level^49^. Furthermore, we propose a concrete microcircuit mechanism for dendritic gating that is constrained by empiricalfindings and makes a number of testable predictions.

The gating mechanism as studied here is nonlinear but not binary. In the biologically plausible regime of inhibitory strength studied here, shunting inhibition on a dendritic branch still allows synaptic input to appreciably elevate the dendritic voltage and thus impact the soma, which decreases the gating selectivity of the neuron. Gating selectivity is also limited by the number of dendritic branches (or more generally, quasi-independent computational units) on a pyramidal neuron. Due to these limitations, our mechanism may be better suited to coarse gating of distinct pathways rather than transmission of more fine-grained top-down signals. Multiple mechanisms may jointly contribute to gating function, and our proposed mechanism is most likely compatible with the aforementioned proposals.

## Supplemental Information

Supplemental Information includes eight figures, two tables and a Supplemental Mathematical Appendix.

## Author Contributions

G.R.Y., J.D.M., and X.-J.W. designed the research, discussed regularly throughout the project and wrote the manuscript. G.R.Y. performed the research.

## Acknowledgments

We thank Michael Higley, Alex Kwan, Jorge Jaramillo, and Francis Song for comments on an earlier version of the manuscript. Funding was provided by National Institute of Health grant R01MH062349 and Office of Naval Research grant N00014-13-1-0297 (X.-J.W.).

## Online Methods

A summary of all types of models used and where they are used can be found in **Supplementary Table 2**.

### Spiking pyramidal neuron models

For the fully reconstructed multi-compartmental pyramidal neuron model (**Supplementary Fig. 1a-d**),weadapted apreviously developed model based onalayer 2/3 pyramidal neuron inthe rat somatosensory cortex reportedby [51]. We used the passive membrane parameter set; results are essentially the same with the active membrane parameter set. Simulations were implemented with the NEURON simulator^52^.

The reduced multi-compartmental spiking neuron model is comprised of multiple dendritic compartments and one somatic compartment. All dendritic compartments are equivalent, not directly coupled to each other, and coupled to the soma. There are 10 dendritic compartments for all simulations using this model (**Fig. 2**,**7**). The number of dendrites does not change the results as long as we normalize the dendrite-soma coupling strength with respect to the number of dendrites. The soma is modeled as a leaky-integrate-and-fire compartment with dynamics following:

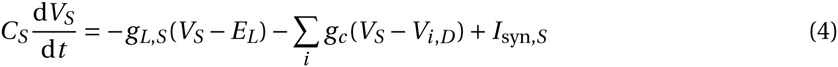

where the subscripts *S* and *D* correspond to soma and dendrites, respectively. *V*_*i,D*_ is the membrane potential of the *i*-th dendrite. *C*_*S*_ is the membrane capacitance, *E*_*L*_ is the resting potential, *g*_*L*_ is the leak conductance, *g*_*c*_ is the coupling between each dendritic compartment and the somatic compartment. We set *C*_*S*_ = 50.0 pF and *g*_*L,S*_ = 2.5 nS, producing a 20-ms membrane time constant for soma. We also set *E*_*L*_ = −70 mV and *g*_*c*_ = 4.0 nS. The somatic spiking mechanism is integrate-and-fire, with spike threshold −50 mV, reset potential −55 mV, and refractory period 2 ms. The dynamics of the dendritic membrane potential (*V*_*D*_) follows

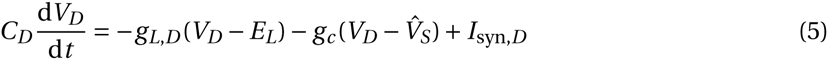

where 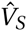 is the somatic shadow potential^53^, which follows the same equation as *V*_*S*_, except with no spiking and resetting. We set *C*_*D*_ = 20.0 pF and *g*_*L,D*_ = 4.0 nS, producing a 5-ms membrane time constant^54^. After a somatic spike, the back-propagating action potential is modeled as an 3-ms-delayed voltage increase of 10 mV in all dendrites^55^.

The main free parameters of the reduced-compartmental model, *g*_*c*_ and *g*_*L,D*_, were chosen to match *in vitro* properties reported by [54]. Specifically, a single-synapse dendritic EPSP of 1-mV peak is attenuated to about 0.05mVinthe soma, and a dendritic NMDA plateau potential evokes a somatic depolarization with the peak around 10 mV. We also made several efforts to adapt our model to mimic physiological *in vivo* conditions, including excitation-inhibition balanced background inputs and reduced soma-dendrite coupling. We used an *in-vivo* set of parameters whenever appropriate (**Fig. 2d-g** and **Fig. 7c,d**). The soma-dendrite coupling is reduced fivefold to *g*_*c,vivo*_ = 0.8 nS, to achieve the stronger signal attenuation observed in high-conductance state^56^. In this regime, the soma also receives excitatory and inhibitory background inputs, 500 Hz of 2.5-nS AMPAR input and 150 Hz of 4.0-nS GABAR input, to approximate the excitation-inhibition balanced background input that gives the neuron a baseline Poisson-like firing rate around 3 Hz. Reduced spiking neuron simulations were implemented with the BRIAN neural simulator^57^. All simulation codes are available on ModelDB.

We used four types of synapses, AMPAR, NMDAR, GABA_A_R, and GABA_B_R. Since GABA_B_Rs are only used briefly in (**Supplementary Fig. 3**), we denote GABA_A_ simply as GABA. AMPAR and GABAR synapses are modeled as linear:

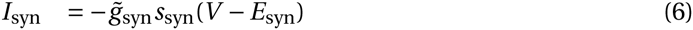

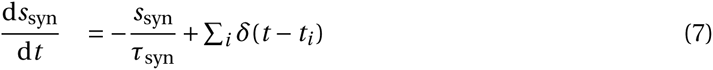

where *s*_syn_ is the gating variable representing the proportion of open channels, 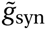 is the maximum synaptic conductance, *E*_syn_ is the synaptic reversal potential, *τ*_syn_ is the synaptic time constant, and *t*_*i*_ are pre-synaptic spike times. We set *τ*_AMPA_ = 2 ms, *E*_AMPA_ = *E*_*E*_ = 0 mV, *E*_GABA_ = *E*_*I*_ = −70 mV, and 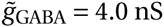. For dendrite-targeting inhibitory synapses *τ*_GABA,dend_ = 20 ms, whereas *τ*_GABA,soma_ = 10 ms for soma-targeting inhibitory synapses. These are based on the observations that dendrite-targeting inhibition tend to be slower^58,59^. In **Supplementary Fig. 1d,h**, 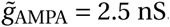 In **Supplementary Fig. 3**, 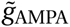 ranges from 0 to 2.5 nS. Otherwise 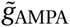 is set as 0 nS (no AMPAR input).

GABA_B_R synapses are post-synaptic. Each spikeat time *t*_*i*_ increases the gating variable *s*_GABA_B__(*t*) by *γ*_GABA_B__[exp[*t* − *t*_*i*_)/*τ*_GABA_B_,decay_] − exp [*t* − *t*_*i*_)/*τ*_GABA_B_,rise_]], where *γ*_GABA_B__ is a normalizing factor such that the peak of the above expression is 1. Then the total input current voltage dependent is

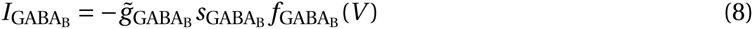

where *ƒ*_GABA_B__(*V*) = 33.33mV · (0.5-2/(1 + exp((*V* + 98.73)/12.5))), as obtained from [60].

NMDAR synapses include a voltage-dependent magnesium block *ƒ*_Mg_(*V*) and saturating gating variable S_NMDA_:

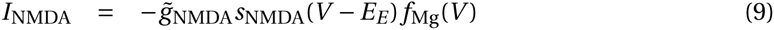

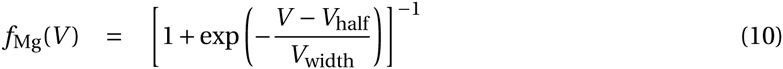

with *V*_half_ = −19-9 mV and *V*_width_ = 12.48 mV^61^. The NMDA conductance 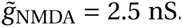. The NMDAR gating variable dynamics follow:

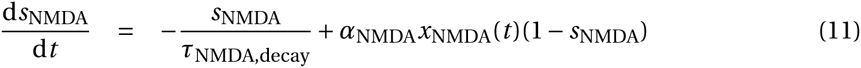

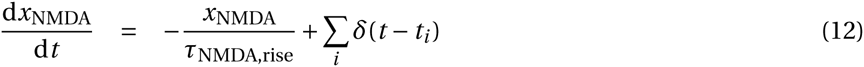

with *τ*_NMDA,decay_ = 100 ^ms^, *τ*_NMDA,rise_ = 2 ms, and *α*_NMDA_ = 0.3 ms^−1^. This choice of *α*_NMDA_ sets SNMDA to be roughly 0.4 at its peak after a single spike^62,63^. With this value of *α*_NMDA_, the saturation of NMDA starts to get prominent around firing rate *r* = 1/(*α*_NMDA_*τ*_NMDA,rise_*τ*_NMDA,decay_) ≈ 16 Hz. By default in simulations with the reduced spiking model, the excitatory inputs are 15 independent NMDAR synapses with the same rate. Fewer number of excitatory synapses can become insufficient to elicit NMDA plateau potential. Since GABAR and AMPAR synapses are linear, their inputs are directly represented by the overall rates.

Each excitatory synapse also has a calcium concentration level with arbitrary unit, which consists of two components, one NMDAR-dependent and one voltage-gated calcium-channel (VGCC) dependent: [Ca^2+^] = [Ca^2+^]_NMDA_ + [Ca^2+^]_VGCC_. The NMDAR-dependent component is modeled as leaky integration of the NMDAR current:

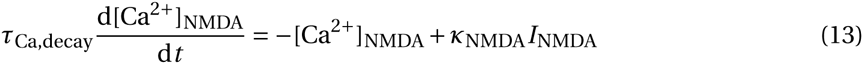

where *κ*_NMDA_ is a scaling parameter with unit pA^−1^. The VGCC component is evoked by postsynaptic spikes that back-propagate into dendrites. Each spike induces a bi-exponential increase:

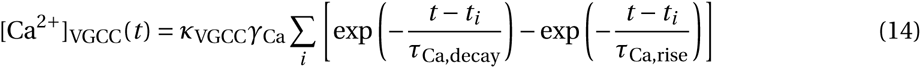

Here *γ*_Ca_ is a normalizing constant so that the peak response to one spike is *κ*_VGCC_. And *κ*_VGCC_ is again a scaling parameter. *τ*_Ca,decay_ = 30 ms is estimated from [35]. *τ*_Ca,rise_ = 2 ms is used mainly to make [Ca^2+^] continuous.

### NMDA plateau potential

The voltage of a dendrite receiving NMDAR and GABAR inputs follows

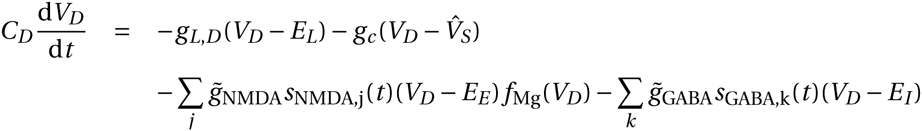

where *j* and *k* are indices of NMDAR and GABAR synapses respectively. Denote

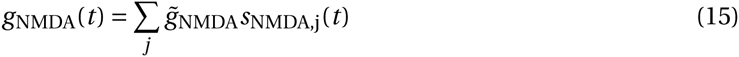

as the total NMDA input conductance onto this dendrite. The maximum value of *g*_NMDA_ (*t*) is simply 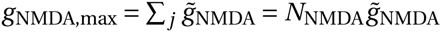, where *N*_NMDA_ is the number of NMDAR synapses. Similarly

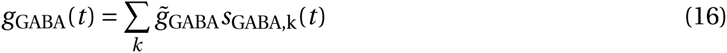

If we ignore the coupling between this dendrite and its soma for now, and consider constant synaptic conductances *g*_NMDA_ = *g*_NMDA_(*t*), *g*_GABA_ = *g*_GABA_(*t*). Then we have

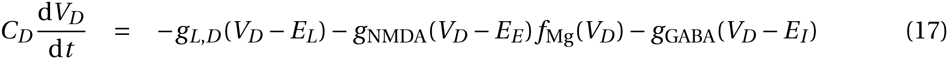

Since we have *E*_*I*_ = *E*_*L*_, the steady-state dendritic voltage *V*_*D*,ss_ satisfies

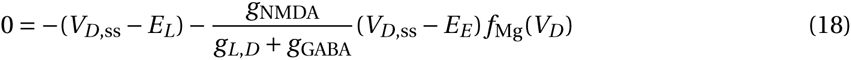

This equation can be solved numerically resulting in the curve in **Supplementary Fig. 2d**.

### Pathway-specific gating in single pyramidal neuron

Gating is performed by disinhibiting a specific subset of dendrites. Disinhibited dendrites always receive 5 Hz background inhibition. The disinhibition level is defined as the difference between the rates of inhibition received by inhibited and disinhibited dendrites.

In **Fig. 2d-g**, each pathway targets two dendrites with 15 NMDAR synapses on each dendrite. The dendrites targeted by each pathway do not overlap. For each pathway the input rate (*u*_*E*_) follows a bell-shaped tuning to the stimulus value (*z*): *u*_*E*_ = 40exp(−*z*^2^) Hz, where *z* ranges between −2.4 and 2.4. The disinhibition level is 30 Hz (from 35 Hz to 5 Hz).

Presented alone, the preferred stimulus (*z* = 0) from one pathway increases the output firing rate by *r*_on_ (*r*_off_) when the pathway is gated on (off). The gating selectivity is defined as

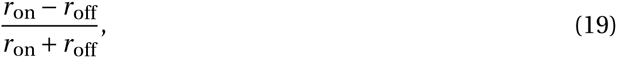

For **Fig. 3**, excitatory pathways can overlap. In the context with gate 1 open, *N*_disinh_ dendrites are disinhibited. Excitatory pathway 1 targets these *N*_disinh_ dendrites, each with strength 25 nS, and similarly for gate 2 and pathway 2. The *N*_disinh_ dendrites disinhibited for gate 2 are chosen randomly and independently from the *N*_disinh_ dendrites disinhibited for gate 1. For each *N*_disinh_ and *N*_dend_, *r*_on_, *r*_off_ are averaged across all possible projection patterns.

### Rate pyramidal neuron model

The rate model is fitted with simulation data from the spiking model with *in-vivo* parameters (**Supplementary Fig. 4**). The time-averaged voltage of a dendritic compartment 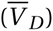 is modeled as a sigmoidal function of total excitatory input conductance (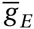, see below for definition) following:

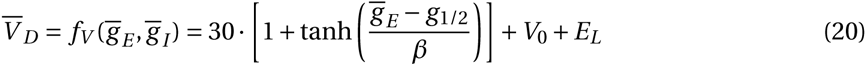

The mid-point *g*_1/2_ is proportional to the total inhibitory conductance 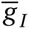 plus the leak conductance of the dendrite *g*_*L,D*_, as expected from the constant conductance scenario **Supplementary Fig. 2c**)

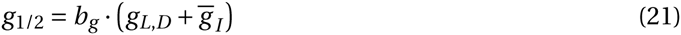

Based on our observation of the reduced spiking model, we modeled the width *β* as an exponentially increasing function of inhibition:

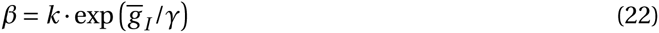

This increase of width *β* as a function of 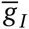 captures the linearization effect of sparse inhibition on the voltage-input function (**Fig. 2c**). Fit values of the parameters are *b*_*g*_ = 5.56, *k* = 9.64 nS, *γ* = 6.54 nS, *V*_0_ = 0.78 mV. The model is fitted to a simulated 10-dendrite spiking neuron model. When simulating dendrites of the spiking model, somatic shadow voltage is clamped at −60 mV, and back-propagating action potential is fixed as a Poisson spike train of 10 Hz. This phenomenological model allows us to interpolate the dendritic voltage for a large range of excitatory and inhibitory inputs very rapidly.

The firing rate of the soma is modeled as a power law function of input current *I*:

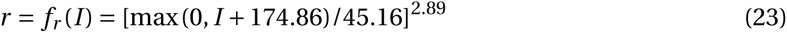

Here *I* is the sum of the input current from dendrites and also the somatic inhibition from PV neurons whenever applicable. The parameters are fitted from simulation of the reduced spiking model. We assume the somatic voltage fluctuates around *E*_reset_ and denote the mean dendritic voltage 〈*V*_*D*_〉. Then the input current from dendrites is *I*_dend→soma_ = *G*_*c*_ · (〈*V*_*D*_〉 − *E*_reset_), where *G*_*c*_ is the total dendrite-soma coupling of all dendrites. *G*_c_ = 8 nS. Since we assume *G*_*c*_ is fixed whenever we vary the number of dendrites (**Fig. 3,4**), the somatic function does not depend on the number of dendrites and need not be re-parametrized. So *I* = *I*_dend→soma_ + Δ*I*_PV→soma_, where Δ*I*_PV→soma_ is the change in somatic inhibition from PV neurons.

For inputs to the rate model, 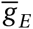 and 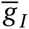 are the time-averaged total conductance of all excitatory and inhibitory synapses, respectively. For NMDAR-only excitatory input, the approximated time-averaged gating variable 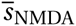 of a single synapse receiving input rate *r*_*E*_ follows,

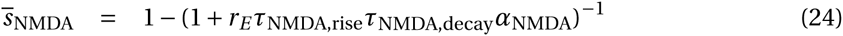

For *N*_NMDA_ synapses each with maximal conductance 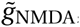, the total excitatory conductance is

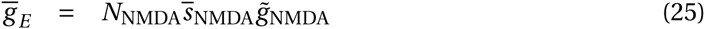

Therefore, 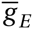 saturates as 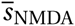 does. Because the GABAR conductance is linear in its input rate, the total inhibitory conductance is

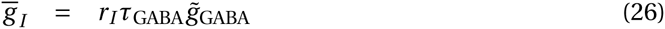

where *r*_*I*_ is the overall inhibitory input rate onto that dendrite.

### Interneuron Models

SOM neurons are modeled as simple rate neurons with a rectified linear f-I curve. The firing rate of a SOM neuron is

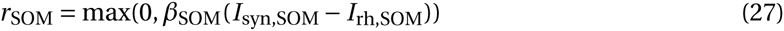

where max(*x*, 0) is a rectified linear function of *x. I*_rh,SOM_ = 40 pA is the rheobase, i.e. the minimum current required to activate the neuron, and *β*_SOM_ = 90 Hz/nA is the f-I curve slope for SOM neurons, which we matched to data from [64]. SOM neurons typically display adapting responses to constant input, and the synapses of SOM neurons show short-term-plasticity We ignored these aspects of temporal dynamics because here we are interested in the steady-state response. SOM neurons receive 150 pA input current in the default state, leading to a baseline firing of SOM neurons around 10 Hz as observed experimentally^5,13^.

For VIP neurons, we assume that the control input targets *N*_control,VIP_ = round(*N*_VIP_·*P*_control,VIP_) of them. On average VIP neurons are assumed to fire at 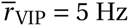. Therefore the VIP neurons non-activated by the control input fire at 0 Hz, while those targeted by the control input fire at (5 · *N*_VIP_/*N*_control,VIP_)Hz.

PV neurons are modeled simply as linear rate neurons with a f-I curve slope of *β*_PV_ = 220 Hz/nA, because their activities never reach zero in our model. Since we are only interested in their change of activities in response to SOM neuron suppression, the spontaneous activity of PV neurons is irrelevant to our model.

### Interneuronal Network

The full interneuronal network model contains pyramidal, SOM, VIP, and PV neurons. The network model is roughly based on a L2/3 cortical column microcircuit, and contains 3000 pyramidal neurons, 160 SOM neurons, 140 VIP neurons, and 200 PV neurons^65^. However, the analysis applies more generally. Pyramidal neurons are modeled as multi-compartmental rate neurons as described above. We typically used *N*_dend_ = 30 dendrites, approximately corresponding to pyramidal neurons in associative areas^66^. The number of pyramidal neurons does not affect our results.

The SOM-to-dendrite connections are set randomly. Instead of drawing each connection randomly and independently with a fixed probability we assume that each dendrite is targeted by precisely *N*_SOM→dend_ SOM neurons, when *N*_SOM→dend_ is an integer, so that each dendrite receives the same amount of net inhibition in the default state. The identities of SOM neurons targeting each dendrite is randomly chosen. The total inhibitory conductance received by each dendrite is denoted and fixed as *G*_SOM→dend_ = 40 nS, then for each SOM-dendrite connection the conductance is *G*_SOM→dend_/*N*_SOM→dend_. Each SOM-dendrite connection can contain multiple synapses, which we are not explicitly modeling here because GABAergic synapses are linear such that we only need to consider the total conductance. When *N*_SOM→dend_ is not an integer, we interpolate. Each dendrite is targeted by ⌈*N*_SOM→dend_⌉ SOM neurons, where all synapses but one have weights *G*_SOM→dend_/*N*_SOM→dend_, while one has weight *G*_SOM→dend_ · (1 -⌊*N*_SOM→dend_⌋ /*N*_SOM→dend_). Given the connection probability from SOM to pyramidal neurons *P*_SOM→pyr_, the number of SOM neurons *N*_SOM_, and the number of dendrites on each pyramidal neuron *N*_dend_, we set

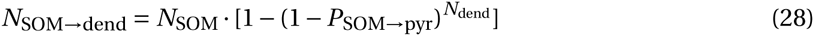

This is the mean number of SOM neurons targeting each dendrite if the SOM-to-pyramidal connections were completely independent and random.

The VIP-to-SOM connections are set in the same way as the SOM-to-dendrite connections. Since SOM neurons only have one compartment each, we have *N*_VIP→SOM_ = *N*_VIP_ · *P*_VIP→SOM_. When control inputs target both VIP and SOM neurons, we have *P*_VIP→SOM_ = 0.6. When control inputs only target VIP neurons, we have *P*_VIP→SOM_ = 0.1. Within 100//m the connection probability is measured to be around 0.6 (**Supplementary Table 1**). However, note that the connection probability from VIP to SOM neurons on a column scale is unknown. The spatially-restricted ax-onal arbors of VIP neurons^31^ suggest that the connection probability may fall quickly as a function of the VIP-SOM distance. Therefore on the scale of a column, the connection probability could still be as low as 0.1. Total inhibitory connection weight from VIP neurons received by each SOM neuron is 30 pA/Hz, and is distributed onto each synapse the same way SOM-to-dendrite connection weights are set. For *N*_VIP_ = 140 and *P*_VIP→SOM_ = 0.6, the connection strength of each synapse is about 0.4 pA/Hz. This is close to the unitary VIP-to-SOM IPSQ of 0.7 pA/Hz measured in [4]. Notice here the connection is current-based because SOM neurons are described by a f-I curve.

The SOM-to-PV, PV-to-PV, and PV-to-pyramidal soma connections are all set similar to the connections above. We set *P*_SOM→PV_ = 0.8, *P*_PV→PV_ = 0.9, *P*_PV→soma_ = 0.6^4^. The total inhibitory connection strength from SOM neurons to each PV neuron is varied in **Fig. 6b**. The total inhibitory connection from PV neurons to each PV neuron is 30 pA/Hz, and from PV neurons to each pyramidal neuron is 30 pA/Hz. Denote the resulting connection weight matrix *W*_SOM→PV_, *W*_PV→PV_, *W*_PV→soma_, then in steady state the change in somatic inhibition Δ**I**_pyr_ across pyramidal neurons is

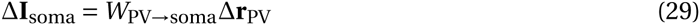

where Δ**r**_PV_ is the change in PV activities. And we have

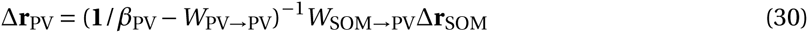

where Δ**r**_SOM_ is the change in SOM activities before and after control inputs. **1** is identity matrix. The precise values of these PV-related parameters do not matter.

Control inputs are excitatory. Here we are agnostic about their origin. They could be locally generated or from long-range projections. Control inputs can target subsets of SOM and VIP neurons. The mean strength of the control inputs across the whole population is always kept as a constant. When control inputs target SOM neurons, *N*_control,SOM_ = round(*N*_SOM_ · *P*_control,SOM_) of SOM neurons are targeted, with current 75·*N*_SOM_/*N*_control,SOM_ pA. Therefore across the whole population the averaged current input is 75 pA. When control inputs target VIP neurons, each of the *N*_control,VIP_ targeted VIP neurons fire at (5·*N*_VIP_/*N*_control,VIP_) Hz. For **Fig. 5a-c** when control inputs only target VIP neurons, we set *P*_control,SOM_ = 0,*P*_control,VIP_ = 0.1. *P*_control,VIP_ has to be low otherwise the gating selectivity would be very close to 0. For **Fig. 5d-f**, when control inputs target both SOM and VIP neurons, *P*_control,SOM_ = 0.5,*P*_control,VIP_ = 0.5. Setting *P*_control,SOM_ = 0.5 does not result in the highest gating selectivity. We did not make particular efforts to fine-tune these parameters.

Finally, excitatory inputs carrying stimulus information for one pathway target the corresponding disinhibited dendrites. When we opened the gate for pathway 1, suppose one dendrite receives averaged inhibitory conductance *g*_*I*_. Then the total excitatory conductance 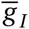 from pathway 1 onto this dendrite is

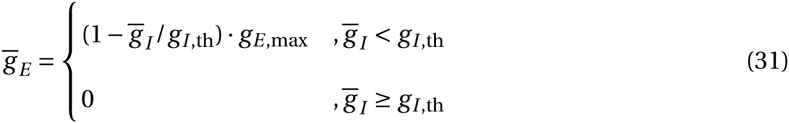

*g*_*I*,th_ is a inhibitory conductance threshold we defined. Therefore when inhibition is weak (disinhibition is strong), excitation is inversely proportional to inhibition level. However, when disin-hibition is weak, there will be no excitatory input at all. Having a cut-off threshold *g*_*I*,th_ prevents excitatory inputs from targeting every dendrite and therefore being overly dense. We set *g*_*I*,th_ = 4.0 nS. Since we have set the sum of conductances of all inhibitory synapses to be *G*_SOM→dend_ = 40 nS, each SOM neuron fires around 10 Hz prior to disinhibition, and *τ*_GABA,dend_ = 20 ms, the time-averaged conductance received by each dendrite in default is 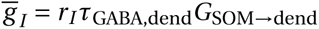 = 8.0 nS. Therefore by setting *g*_*I*,th_ = 4.0 nS, excitatory synapses only target dendrites that are at least disinhibited by half. We set the maximum time-averaged excitatory conductance targeting each dendrite to be *g*_*E*,max_ = 25 nS. This value is chosen so that excitation is strong enough to excite a disinhibited dendrite, but not strong enough to excite a strongly inhibited dendrite (**Supplementary Fig. 4**).

In (**Supplementary Table 1**), we summarized the raw experimental data used to constrain the model.

### Synaptic plasticity model and learning protocol

The synaptic plasticity model is calcium-based. The calcium dynamics is described above, and the synaptic weight change given these calcium dynamics is modeled with the formalism from [34], restated below for clarity with slightly modified notations.

Over the time of stimulation, the calcium trace spends time *α*_*p*_ above the potentiation threshold *θ*_*p*_, and time *α*_*d*_ above the depression threshold *θ*_*d*_. Then the average potentiation is Γ_*p*_ =*γ*_*p*_*α*_*p*_, and the average depression is Γ_*d*_ =*γ*_*d*_*α*_*d*_, where *γ*_*p*_ and *γ*_*d*_ are the potentiation and depression strengthes respectively. Since the synapse is assumed to be bistable (DOWN or UP states), denote *ρ* as the probability of the synapse staying in the UP state, which evolves over time in response to the calcium trace crossing thresholds. Then define 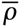 as the long-term time average of *ρ*, and 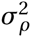 as the standard deviation of *ρ*. Then

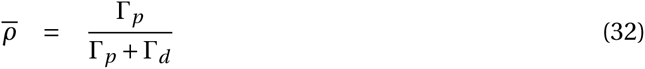

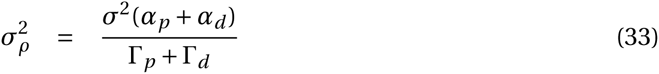

where σ is the amplitude of noise and *τ* is the time constant of weight change. In long term, the probability to switch from DOWN to UP state 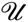 and from UP to DOWN states 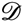 are given by

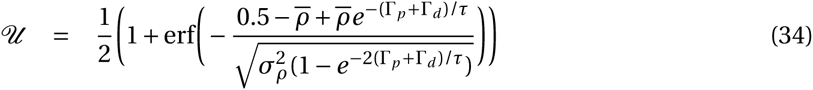

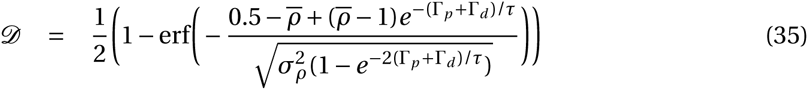

erf(·) is the standard error function. For convenience, we set the weight of DOWN state to *w*_0_ = 0, and the weight of UP state *w*_1_ = 3. Then following stimulation, the weight after learning *w*_post_ = 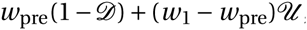, given the weight before learning *w*_*pre*_.

We fitted the free parameters of the model with experimental data from [35]. In simulation of the plasticity experiment, we modeled the pre-synaptic extracellular stimulation by 40 NMDAR synapses simultaneously receiving one spike. This stimulation alone causes a 2.8 mV depolarization on the soma, which is within the range of observed values (1-3 mV) for that experiment. It also brings the dendrite close to the NMDA plateau threshold, allowing for strong interaction between pre-and post-synaptic spikes. As in the experiment, all pairings are repeated 60 times. The somatic shadow voltage is clamped at −60 mV.

The model is fitted to data points from **Fig. 2** **and** **3d** in [35], and is used to predict data from **Fig. 3b** therein. Notice that two data points in the test dataset (their **Fig. 3b**) are already included in their **Fig. 2** **and** **3d**. In all of these cases, there is one pre-synaptic spike, and usually a burst of post-synaptic spikes. The time lag in **Supplementary Fig. 7a** is defined as the timing difference between the first post-synaptic spike in the burst and the pre-synaptic spike. In **Supplementary Fig. 7a,b**, there are 3 post-synaptic spikes. In **Supplementary Fig. 7b,c**, the pre-synaptic spike is either 10 ms earlier than the first post-synaptic spike in the burst, or 10 ms later than the last post-synaptic spike. In **Supplementary Fig. 7a,c**, the post-synaptic burst, when there is one, has frequency of 50 Hz (inter-spike-interval of 20 ms). The fit parameters are the following. The scaling parameters for calcium traces, *κ*_NMDA_ = 0.371 and *κ*_VGCC_ = 0.957. The depression and potentiation rates are *γ*_*d*_ = 39.9 and *γ*_*p*_ = 177.6. The potentiation threshold for calcium is *θ*_*p*_ = 2.78. In fitting this particular dataset, we found that there is a certain level of redundancy in parameters; the number of parameters needed to be free is less than the total number of potentially free parameters. We therefore fixed two parameters using values in [34] which were fitted to another dataset: the amplitude of the noise *σ* = 3.35 and the time constant of synaptic weight change *τ* = 346.36 s. The depression threshold is *θ*_*d*_ = 1. Before the plasticity-inducing experiment, we have *w*_pre_ = 1 which corresponds to 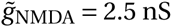 for each NMDAR synapse. So after learning, the actual synaptic conductances are 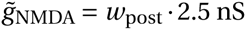.

Just like the spiking pyramidal neuron model, the plasticity model fitted with *in-vitro* data needs to be recalibrated to behave properly under *in-vivo-like* conditions^67^. We reduced the scaling parameters for calcium traces to 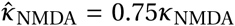, mimicking a reduced extracellular calcium concentration, and to 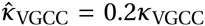, resembling attenuated effect of back-propagating action potentials in high-conductance *in-vivo* state. These changes also ensure the weights of non-activated synapses do not change substantially throughout the simulation. In **Fig. 7b**, the plasticity inducing protocol is 300-s long. The post-synaptic firing is fixed at 10 Hz.

In **Fig. 7c-e**, inputs from both pathways initially target every dendrite with 15 synapses of the same weight 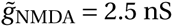. Each gate is opened by disinhibiting 2 distinct dendritic branches. During learning, all synapses from the gated-on pathway are activated at 50 Hz, whereas the gated-off pathway is not activated. The post-synaptic rate is set at 10 Hz. To measure gating selectivity before learning, 8 of the 15 synapses on each dendrite are activated for both pathways. After learning 5 of 15 synapses were activated, the number is chosen so that before and after learning the total excitatory conductance is the same.

### Context-dependent decision-making network

We modeled the context-dependent decision-making task from [26]. In the experimental task, the stimulus is a mixture of random dots that are leftward-or rightward-moving and are red or green. The stimulus can be described by its motion and color coherence. Motion coherence for rightward motion can take 6 values (-0.5, −0.15, −0.05, 0.05, 0.15, 0.5). Color coherence for color red also takes 6 values (-0.5, −0.18, −0.06, 0.06, 0.18, 0.5). On each trial, the color and motion coherence are independently and randomly chosen, resulting in 36 possible stimuli. In **Fig. 8d**, the performance with respect to motion coherence is averaged across all color coherences. Similarly for **Fig. 8e**, the performance with respect to color coherence is averaged across all motion coherences. In **Fig. 8f**, the curve for strong motion coherence is averaged across motion coherence −0.5 and 0.5. Similarly for medium and weak coherences.

The context-dependent decision-making circuit model contains two components. The first is a mixed-selective sensory network, which uses the VIP-SOM-pyramidal neuron circuit model described above. The mixed-selective sensory neurons receive motion and color inputs from the sensory stimulus. Here the motion and color inputs do not signal the actual motion and color of the stimulus, but rather the motion and color evidence for a particular target. For convenience, the motion direction corresponding to target 1 is denoted left, and the color corresponding to target 1 is denoted red, and similar for motion right and color green. This treatment follows the analyses and modeling of [26]. There are four pools of neurons in this network. Each pool prefers a particular combination of motion and color, e.g. left and red. Each neuron pool is modeled exactly as those in **Fig. 5d**, where the circuit connectivity is random and control inputs target both VIP and SOM neurons, using the base parameter set described above. The input to each dendrites is 15 NMDAR synapses with rate determined by the coherence (coh) of their preferred motion and color input, as 40 · (1 + coh) Hz^36,68^. For example, a neuron that prefers left and red inputs would receive 40 · (1 + coh_left_) Hz input on its dendrites targeted by motion pathway, and 40 · (1 + coh_red_) Hz on its dendrites targeted by color pathway. Note that coh_left_ = −coh_right_. The excitatory input for each pathway is set the same way as it is above, however now the maximum conductance of these input synapses *g*_sen_ is one free parameter.

The second component of the network is a decision network. This network is a two-pool rate model^36^, using the parameter set therein with no recurrent AMPAR current. The pool representing choice 1 receives input from a subset of the left-red neuron pool in the mixed-sensory network. Sensory neurons are sorted according to their gating selectivity, and only the top *P*_project_ fraction of these sensory neurons project to the decision networks. *P*_project_ is also a free parameter. The right-green pool projects to the choice 2 pool. The other two pools do not project to the decision network because only the left-red and the right-green pools have congruent preferences for choice 1 and choice 2, respectively, based on the how color and motion evidence are defined. In order to fit experimental behavioral choice data efficiently, we further approximated the decision network with a decision function. We assumed that the probability of selecting choice 1 (*P*_1_) is determined by the difference Δ*I*_dec_ (pA) between the input currents to the two choice pools. We fitted this function by simulating the decision network with mean input current 15.6 pA to both pools, yielding

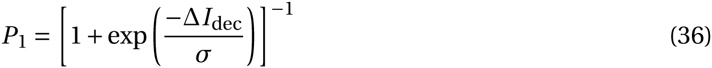

with σ = 0.99 pA. The second free parameter of the model is the projection strength *J*_dec_ of the mixed-sensory input, such that Δ*I*_dec_ = *J*_dec_(*r*_left,red_ − *r*_right,green_). *r*_left,red_ is the average firing rate of the left/red-preferring pool.

The three free parameters *g*_sen_,*P*_project_,*J*_dec_ are fitted to behavioral data of each monkey in [26]. The fit parameter values are *g*_sen_ = 1.21 nS, *P*_project_ = 0.36 and *J*_dec_ = 15.0 pA/Hz for monkey F, and are *g*_sen_ = 1.80 nS, *P*_project_ = 0.083 and *J*_dec_ = 4.37 pA/Hz for monkey A. Importantly, the data used to fit the model is far from being sufficient. Also our circuit model is simplistic. Therefore these fitted parameter values do not reflect our estimates of these quantities in the brain. Rather, these fittings demonstrate that the proposed circuit architecture can potentially capture behavioral performance. As shown in **Supplementary Fig. 8**, if neural gating is strongly degraded, then no set of these fit parameters can capture behavioral performance.

### Model fitting in general

Model parameters are fitted to experimental or simulation data in various contexts. These fitted models include the rate pyramidal neuron, the calcium-based plasticity model, and the context-dependent decision-making network. In all these cases, parameters are chosen to minimize the squared-error between the model and data using sequential least squares programming (SLSQP) method from the SciPy library (scipy.optimize.minimize, with method ‘SLSQP’).

## 1 Supplemental Figures

**Figure S1.**
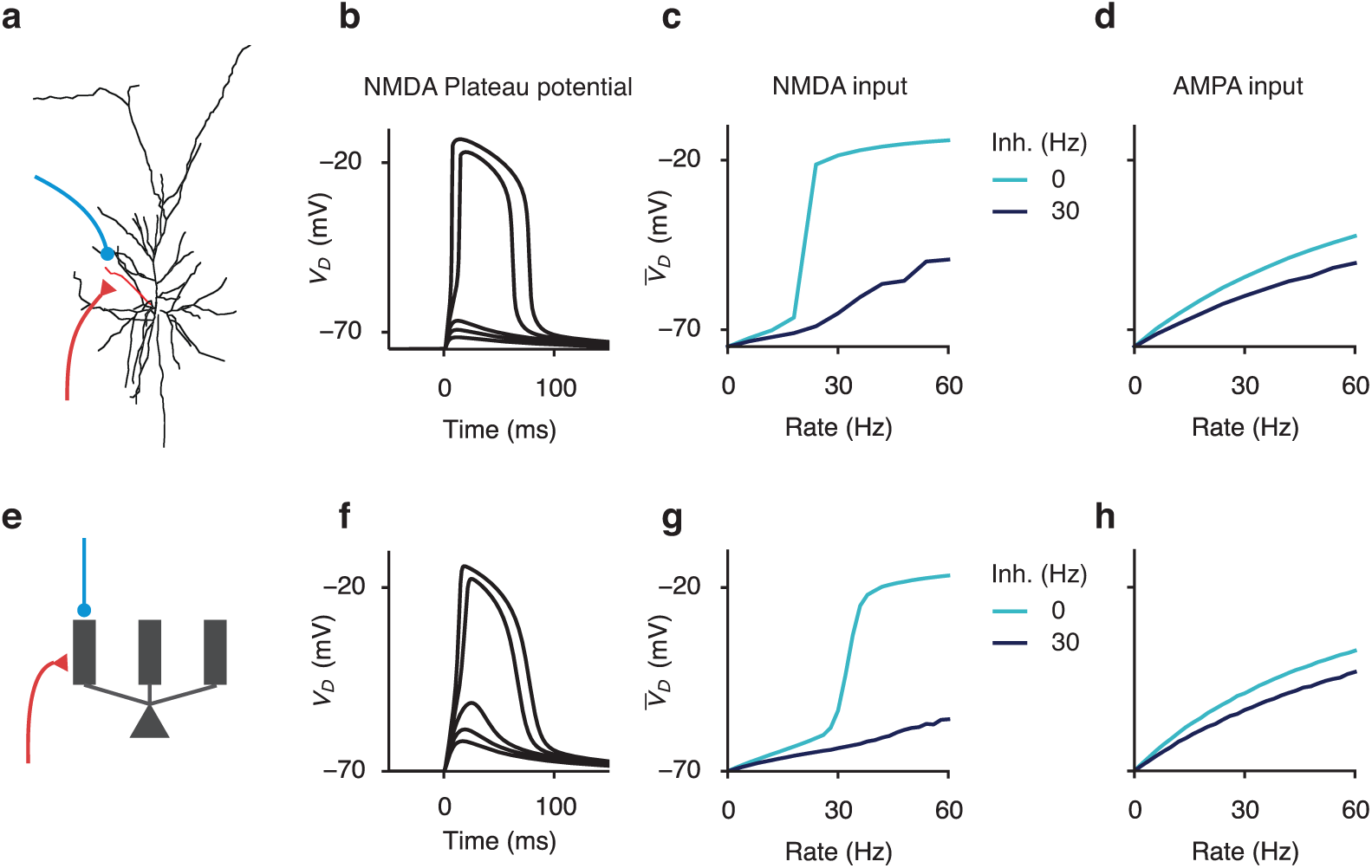
Dendritic disinhibition powerfully gates dendritic nonlinearity. (**a-d**) Dendritic disinhibition controls NMDAR-dependent nonlinearity in a reconstructed compartmental neuron model. (**a**) A morphologically reconstructed compartmental model of a layer 2/3 pyramidal neuron (1) receives excitatory and inhibitory inputs uniformly distributed onto one basal dendrite. (**b**) Excitatory inputs can generate a local, regenerative NMDA plateau potential in the dendrite. As the number of activated synapses is increased, there is a sharp nonlinear increase in the evoked dendritic membrane depolarization (*V*_*D*_). (**c-d**) Presynaptic spike times are modeled as Poisson-distributed events. (**c**) In response to synaptic input mediated by NMDAR channels, the mean dendritic voltage across time 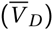 increases nonlinearly as a function of excitatory rate (light blue). Moderate inhibition largely suppresses NMDA plateau potentials even for high excitatory input rate (dark blue). (**d**) The effect of inhibition is much weaker when excitatory input is mediated by AMPARs. 20 excitatory synapses are used as input in (**c,d**). (**e-h**) A reduced compartmental neuron model captures the nonlinearity of the morphologically reconstructed model. (**e**) A somatic compartment is connected to multiple, otherwise independent, dendritic compartments (only three shown). (**f-g**) Modeling results in the reconstructed neuron model (**b-d**) are reproduced by the simplified model. 15 excitatory synapses are used as input in (**g,h**).

**Figure S2.**
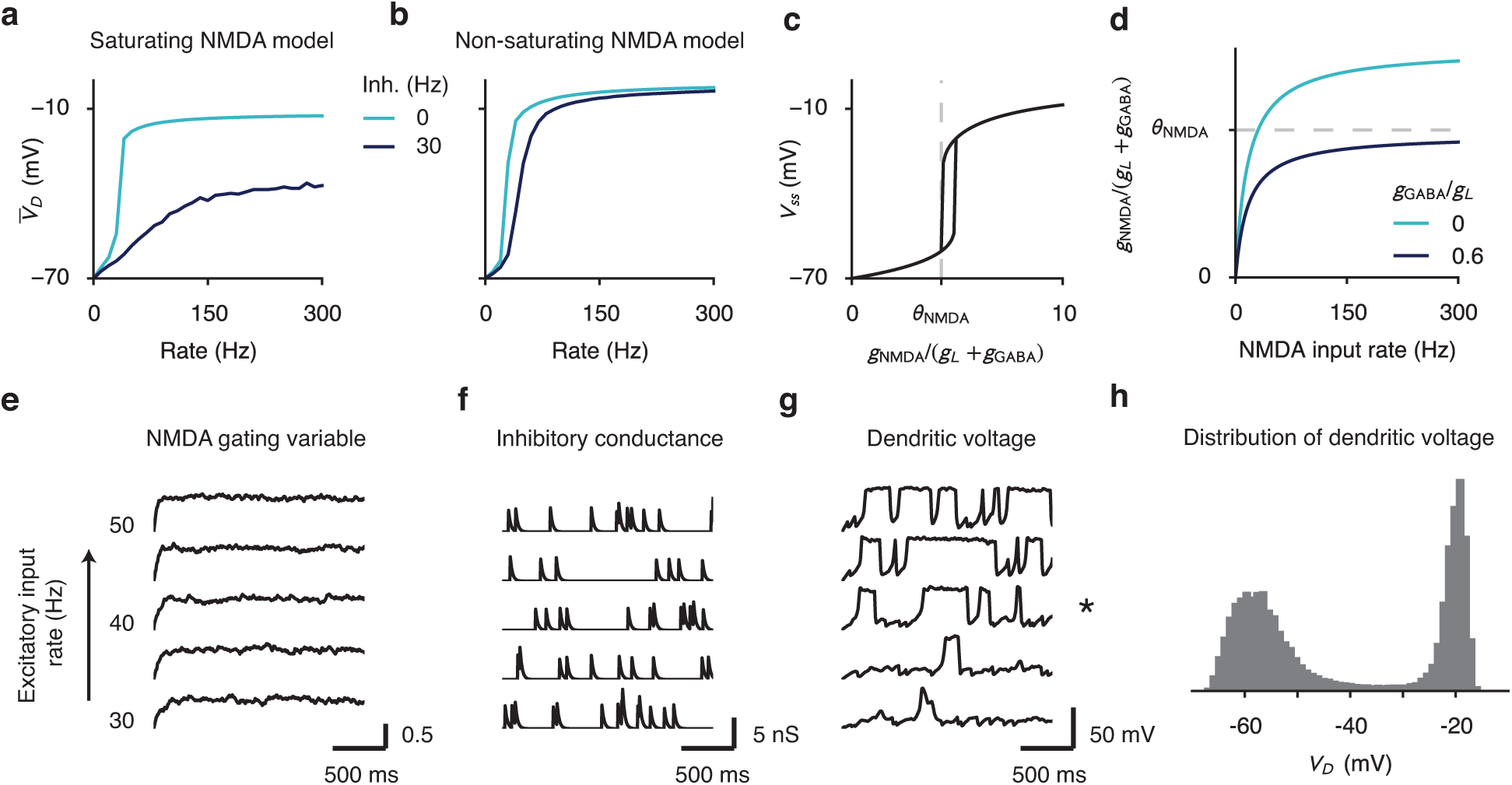
Effects of NMDA receptor saturation and low-rate inhibition. (**a-d**) NMDAR saturation allows for much stronger inhibitory control. (**a**) Using an NMDAR model with saturation allows mild dendritic inhibition to powerfully control dendritic voltage. Note that voltage of the inhibited dendrite (dark blue) never reaches the same level as the disinhibited dendrite (light blue). (**b**) The same level of inhibition has a much smaller effect when we used a non-saturating NMDAR model. (**c**) For constant synaptic conductance, the steady-state voltage of one dendritic branch (*V*_ss_) increases sharply with the effective input *g*_NMDA_/(*g*_*L*_ + *g*_GABA_), where *g*_NMDA_, *g*_*L*_, and *g*_GABA_ are the NMDAR, leak, and GABAR conductances, respectively. The dashed line indicates the threshold *θ*_NMDA_ below which *V*_ss_ is stably in the low state. (**d**) The NMDR conductance, and therefore *g*_NMDA_/(*g*_*L*_ + *g*_GABA_), saturates at high input rates to NMDAR synapses. With moderate inhibition, the saturated value of the effective input can be lower than the threshold *θ*_NMDA_ for an NMDA plateau potential. (**e-h**) Low-rate (temporally sparse) Poisson inhibition generates irregular NMDA plateau potentials and graded encoding of input rate. Inhibition is said to be temporally sparse when the product of the inhibition rate *r*_*I*_ and the time constant *τ*_GABA_ of GABAR is much smaller than 1, i.e. *r*_*I*_ · *τ*_GABA_ ≪ 1 (**e**) Due to relatively high input rate and long time constant, the NMDAR gating variable averaged across synapses is nearly constant in time. Each trace corresponds to a different excitatory input rate, ranging from 30Hz (bottom) to 50Hz (top); the same applies to (**f,g**). (**f**) Inhibitory conductance is temporally sparse due to a low background inhibition rate of 5 Hz. (**g**) The dendritic voltage switches stochastically in time, into and out of the NMDA plateau potential. (**h**) The dendritic voltage across time exhibits a bimodal distribution, due to stochastic switching. The excitatory rate is set to 40 Hz (asterisk in (**g**)).

**Figure S3.**
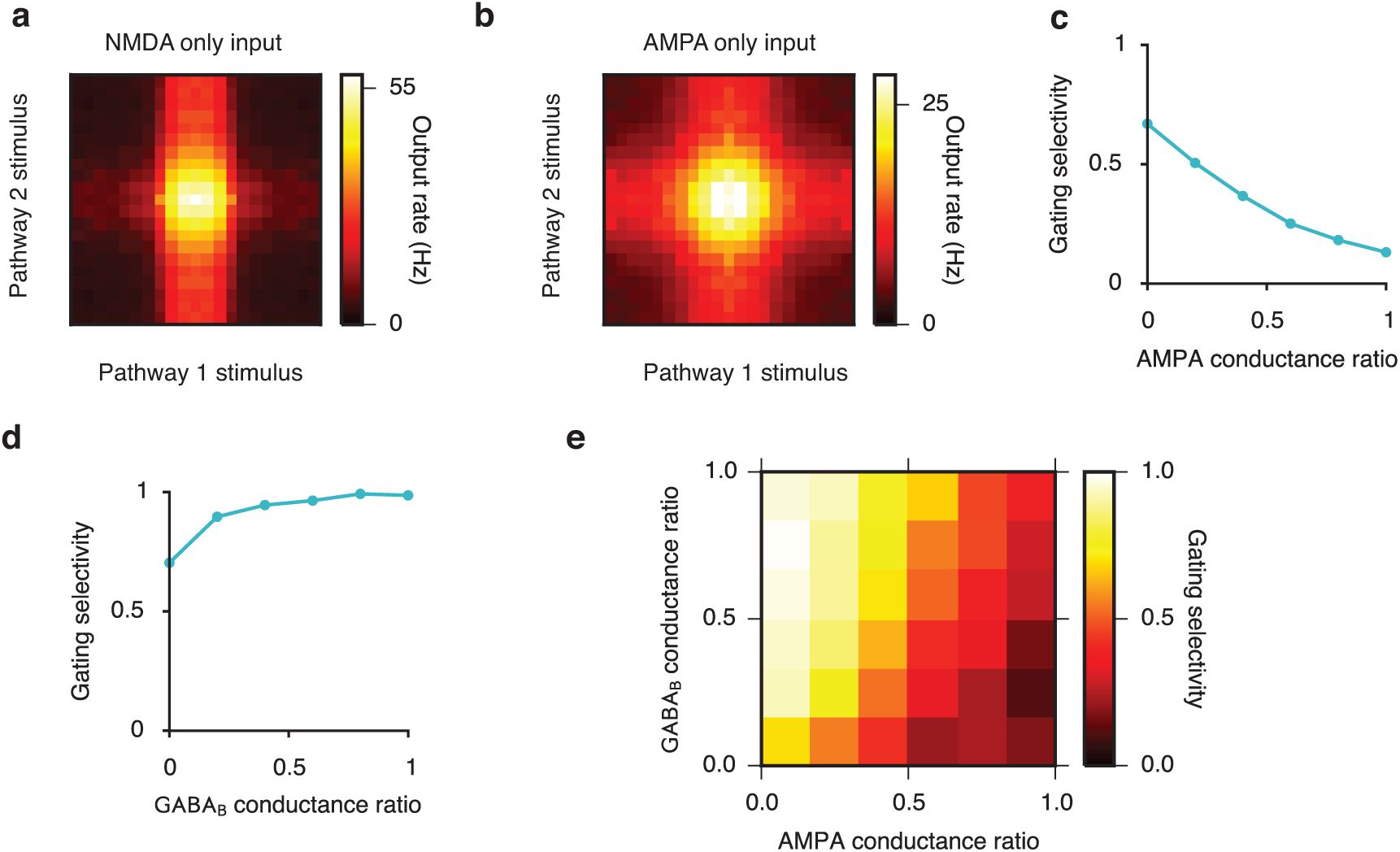
Pathway-specific gating with varying levels of AMPAR and GABA_B_ conductance. In the majority of our work, dendritic excitation is mediated only by NMDARs and dendritic inhibition only by GABA_B_Rs. Here we show how pathway-specific gating varies with the inclusion of AMPAR and GABA_B_ inputs. (**a**) Pathway-specific gating when excitatory input is mediated solely by NMDARs, adapted from Fig. 2g for comparison. (**b**) When the excitatory input is conducted solely by AMPARs (maximum conductance 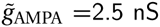 for each synapse), the gating performance is strongly degraded. All other conditions are kept the same in (**a**) and (**b**). Disinhibited dendrites receive 30-Hz disinhibition. (**c**) Gating selectivity (which ranges from 0 for no gating to 1 for perfect gating, see Experimental Procedures for the definition) decreases as a function of the AMPA conductance ratio. Here AMPA conductance ratio is defined as 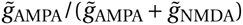, which is 0 in the NMDAR-only case and 1 in the AMPAR-only case. 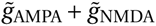 is held constant at 2.5 nS. (**d**) Gating selectivity increases as a function of the GABA_B_ conductance ratio. This is due to both the slower dynamics of GABA_B_ receptors and the inward-rectifying potassium (KIR) conductance activated by GABA_B_ receptors (2; 3). Here excitatory inputs are mediated by NMDARs only. (**e**) Gating selectivity remains high for a wide range of combinations of AMPA and GABA_B_ conductance ratio.

**Figure S4.**
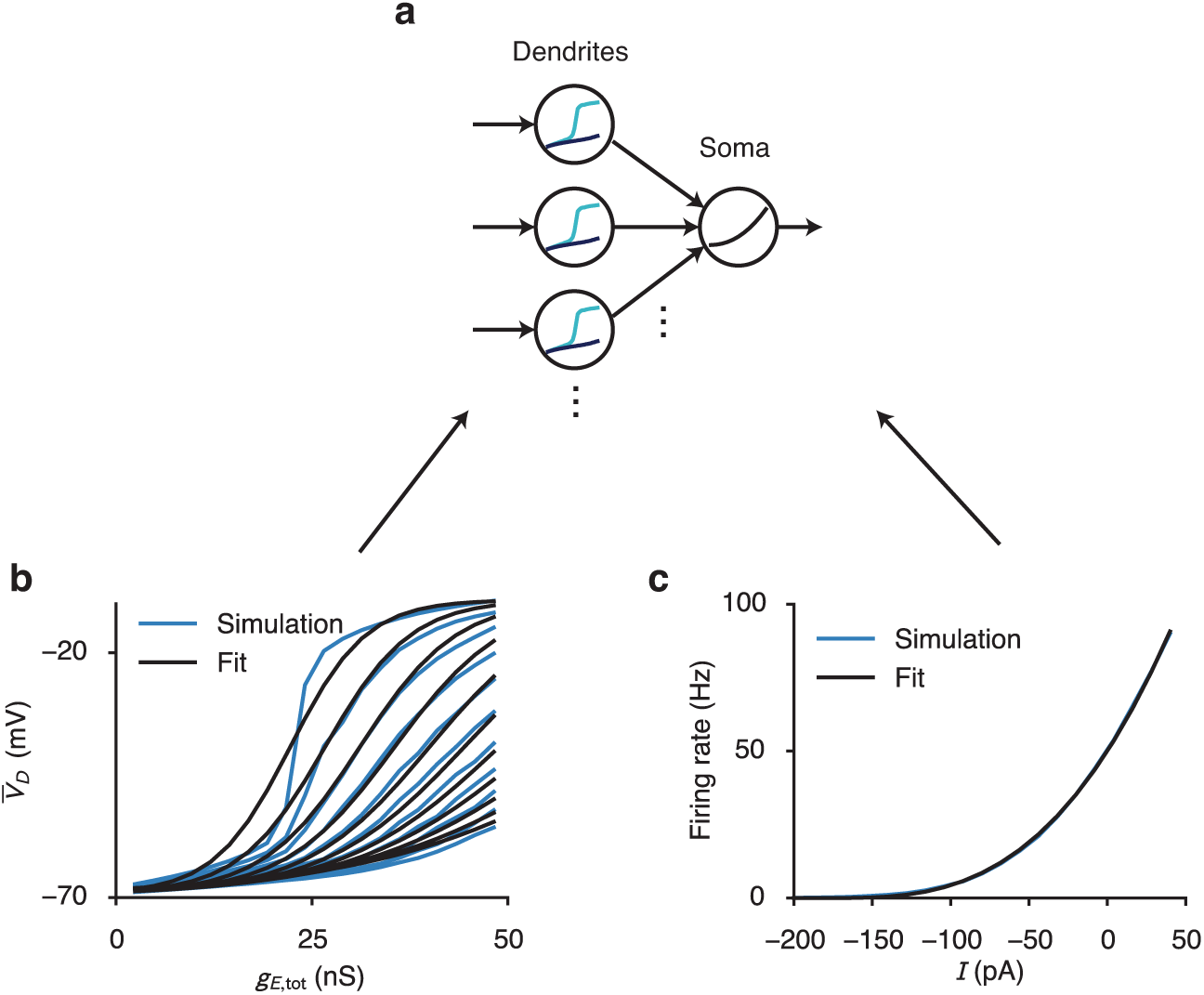
Multi-compartment rate model for pyramidal neurons based on the reduced spiking neuron model. (**a**) The neuron model is comprised of multiple dendrite compartments, whose mean voltages are modeled with a family of sigmoidal functions. These dendritic voltages are converted into currents and fed into a somatic compartment, whose firing rate output is modeled with a power-law function. (**b**) The mean dendritic voltage 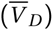 as a function of excitatory and inhibitory inputs. (Blue) Simulation of the reduced-compartmental spiking neuron model. 15 NMDAR inputs fire at a Poisson rate of 30 Hz with conductance ranging from 0.25 to 5.0 nS, resulting in total conductance 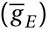 approximately between 0 and 50 nS. Each curve corresponds to a different inhibitory input rate, ranging uniformly from 0 Hz (top curve) to 100 Hz (bottom curve), in increment of 10 Hz. (Black) Fit of the simulation results. All curves are simultaneously fit with a family of sigmoidal functions, where parameters of the sigmoid, i.e. mid-point and width, are controlled by inhibition. The back-propagating action potential is fixed at a rate of 10 Hz. (**c**) Somatic firing rate as a function of input current from dendrites (and potentially PV neurons). In our model, since at resting state the mean dendritic voltage is lower than the somatic voltage, the input current is negative. The simulation result of the spiking model (Blue) is fit with a power-law function (Black).

**Figure S5.**
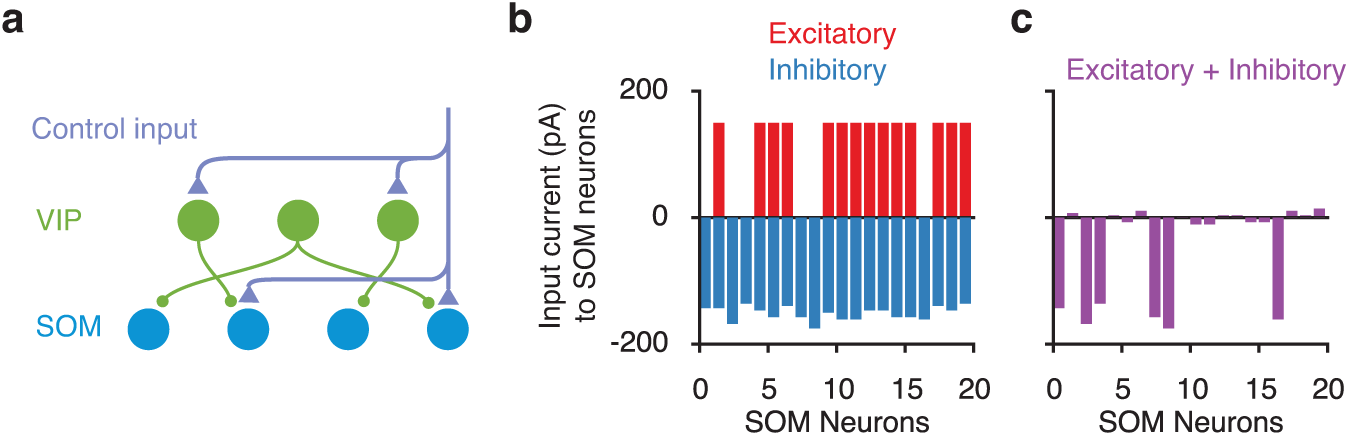
Mechanism of control. **a**) In this scenario, we assume that for each pathway control inputs target a random subset of VIP and SOM neurons. (**b-c**) Input currents onto SOM neurons (only 20 shown). (**b**) 50% of the SOM neurons receive excitatory currents from control (red). 50% of VIP neurons receive excitatory control, but due to the high random connectivity from VIP to SOM neurons, inhibitory currents onto SOM cells are nearly uniform (blue). (**c**) The sum of the excitatory and inhibitory currents onto SOM neurons, i.e. the total currents, are primarily inhibitory and vary strongly across SOM neurons. The overall inhibitory currents are results of overall stronger inhibition. The variability across SOM neurons are mainly inherited from the selective excitatory control input.

**Figure S6.**
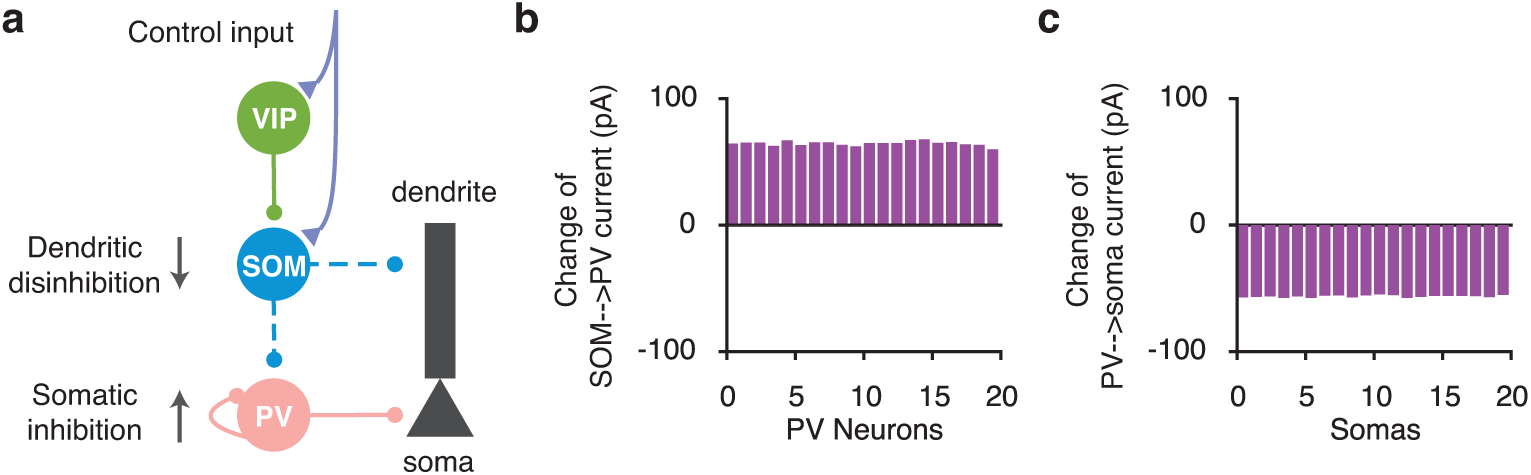
Inclusion of PV neurons results in a uniform somatic inhibition across pyramidal neurons. (**a**) SOM suppression lead to PV disinhibition and somatic inhibition. Same as **Fig. 6a**. (**b**) The change in the SOM-to-PV input currents after the control input. The change in currents is disinhibitory (net excitatory) with a small standard deviation compared to the mean across PV neurons. Notice that although the control input results in a selective suppression of SOM neurons (**Supplementary Fig. 5c**), the change in the SOM-to-PV currents is almost uniform due to the high SOM-to-PV connection probability. (**c**) The change in the PV-to-soma input currents after the control input is net inhibitory and again uniform across somas. Therefore a selective suppression of SOM neurons results in a non-selective inhibition across somas through PV neurons.

**Figure S7.**
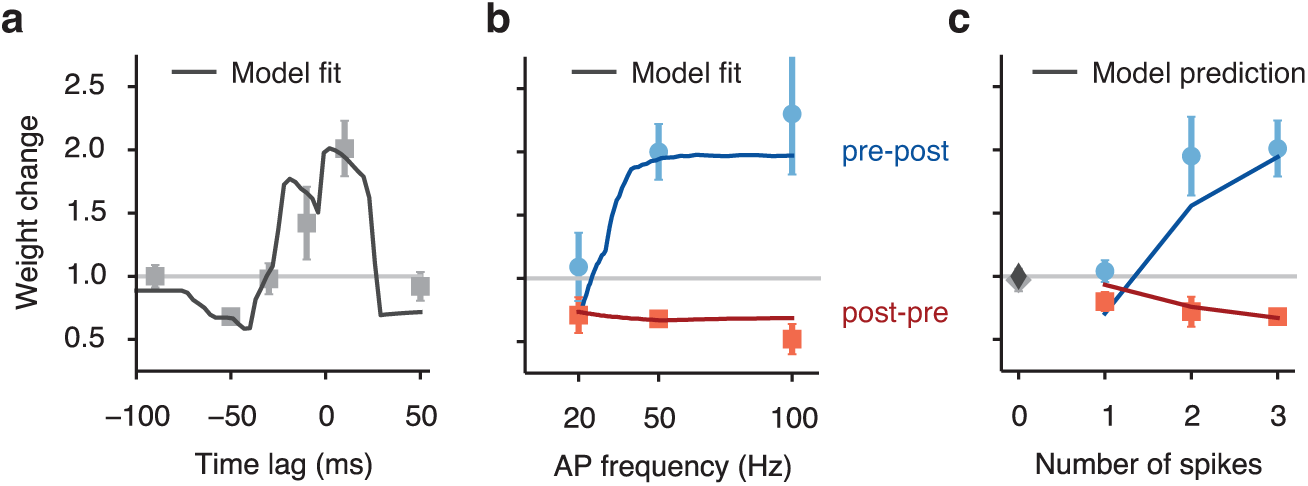
Fit and prediction of the plasticity model compared to experimental data. (**a-c**) A pre-synaptic spike is paired with multiple post-synaptic action potentials (AP). Light symbols mark data showing synaptic weight change (weight after learning/weight before learning) when varying the pre-post time lag (**a**), post-synaptic AP frequency (**b**), and number of post-synaptic spikes (**c**). In (**b,c**), the presynaptic spike either precedes (blue) or follows (red) the postsynaptic spikes. Curves in (**a-b**) show the model fit, with the same set of parameters. (**c**) The model generalizes to predict data not used to fit the model. Experimental data are extracted from (4).

**Figure S8.**
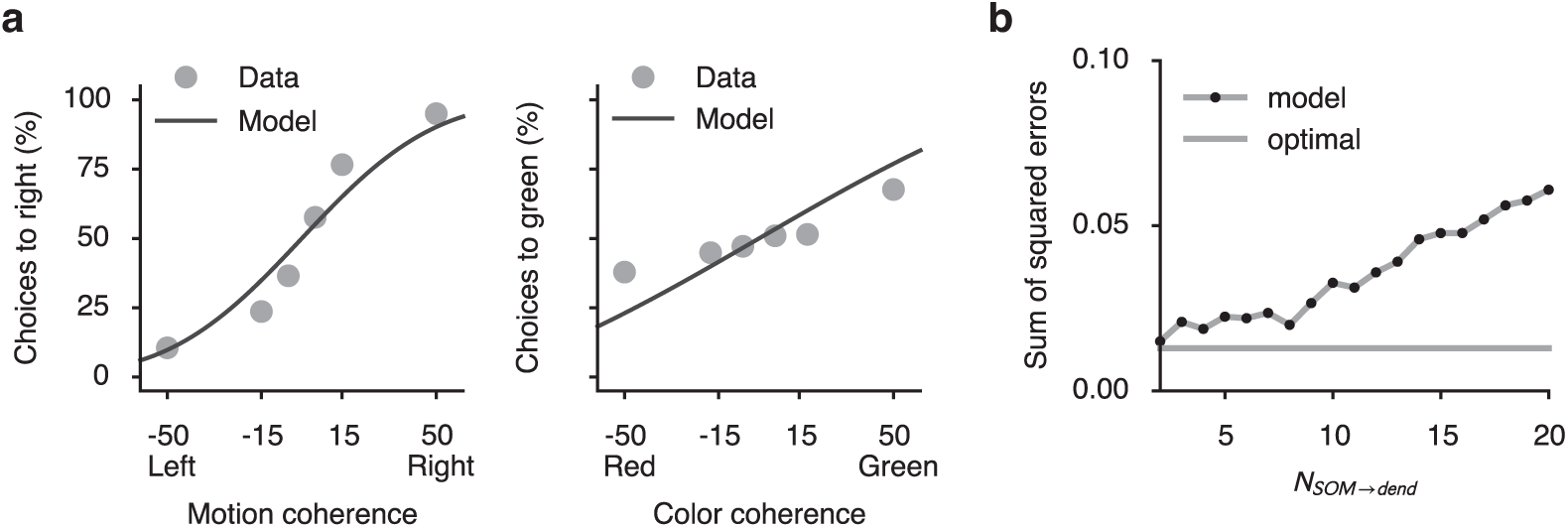
Fits of behavioral data as we vary parameters of the interneuronal circuit. (**a**) The model fit to behavioral data in motion context when we set *N*_SOM→dend_ = 20. The fit is much degraded compared to **Fig. 8**. From **Fig. 4** we know that *N*_SOM→dend_ is the critical parameter for gating selectivity measured on the neural level. (**b**) The sum of squared errors of the model fit as a function of *N*_SOM→dend_. For a large range of *N*_SOM→dend_, the model can nearly fit the data optimally. The fit starts to degrade when *N*_SOM→dend_ > 10. Dashed line indicates the error level of the optimal sigmoidal fit, where data are directly fitted to logistic functions. The sum of squared errors shown here is the median error of 50 different model realizations and fits.

## 2 Supplemental Table

## 3 Mathematical Appendix

### 3.1 Gating selectivity critically depends on *N*_SOM→dend_

The gating selectivity is defined as the mean gating selectivity across neurons,

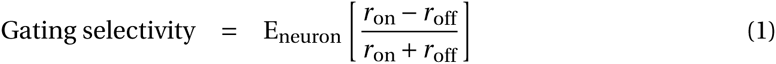

For each neuron, the neural activity given the gated-on pathway is

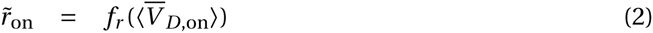
where 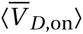 is the mean dendritic voltage across all the dendrites on that neuron for the gated-on pathway. Notice here for simplicity we used an input-output formulation for the somatic compartment that is slightly different from the one used in the main text (the results are the same)

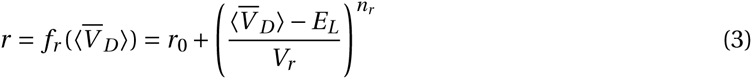

After correcting for the baseline, we have

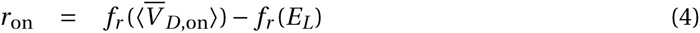

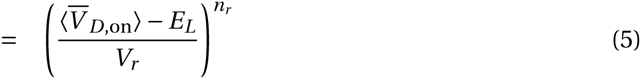

Similarly

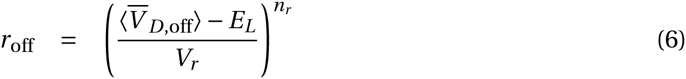

So

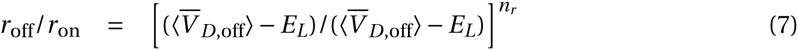

In the limit of large number of dendrites on each pyramidal neuron, we can replace the averaged dendritic voltage with its expectation over dendrites E_D_[·].

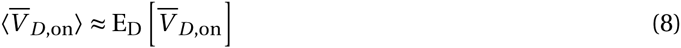

**Table S1.** Raw experimental data used to constrain the VIP-SOM-pyramidal disinhibitory circuit. The error estimates are also taken from the references when available. Some of the data are extracted from their figures since the value is not reported in texts. Specifically, the proportion of VIP neurons is inferred from the proportion of 5HT3a neurons among interneurons and proportion of VIP neurons among 5HT3a neurons.

| Parameter | Value | References | Layer | Area | Animal |
| --- | --- | --- | --- | --- | --- |
| Proportion of SOM neurons among inhibitory neurons | 0.208 | (5) | L2/3 | S1 | mouse |
| Proportion of VIP neurons among inhibitory neurons | 0.2 | (5) | L2/3 | S1 | mouse |
| Total number of inhibitory neurons in a column | $676 \pm 116$ | (6) | L2/3 | S1 | rat |
| Baseline activity of SOM neurons | $6.2 \pm 0.7$ Hz | (7) | L2/3 | S1 | mouse |
| Unitary IPSQ from SOM to pyramidal neurons | $1.5 \pm 0.3$ pC | (8) | L2/3 | V1 | mouse |
| Unitary IPSQ from VIP to SOM neurons | $0.69 \pm 0.33$ pC | (8) | L2/3 | V1 | mouse |
| Connection probability from SOM to pyramidal neurons (within $200\mu\text{m}$) | $0.71 \pm 0.03$ | (9) | L2/3 | frontal cortex | mouse |
| Connection probability from VIP to SOM neurons (within $25 - 100\mu\text{m}$) | $0.625 \pm 0.12$ | (8) | L2/3 | V1 | mouse |
| Number of basal dendrites on each pyramidal cell (number of total tips) | $28.8 \pm 2.4$ | (10) | L2/3 | V1 | rat |
| Number of basal dendrites on each pyramidal cell (maximum branches at fixed radius) | $20 \pm 2.6$ | (11) | L3 | V1 | monkey |
| Number of basal dendrites on each pyramidal cell (maximum branches at fixed radius) | $34.2 \pm 4.9$ | (11) | L3 | anterior cingulate cortex | monkey |

**Table S2.** All types of models used, and their corresponding result figures.

| Model | Figure |
| --- | --- |
| Fully-reconstructed spiking pyramidal neuron model | **Supplementary Fig. 1a-d** |
| Reduced multi-compartmental spiking pyramidal neuron model | **Fig. 2,7, Supplementary Fig. 1e-h,2,3** |
| Multi-compartmental rate pyramidal neuron model | **Fig. 3-6,8, Supplementary Fig. 4-6,8** |
| Rate SOM neurons | **Fig. 4-6,8, Supplementary Fig. 5,6,8** |
| Rate VIP neurons | **Fig. 5,6,8, Supplementary Fig. 5,6,8** |
| Rate PV neurons | **Fig. 6, Supplementary Fig. 6** |
| Calcium-based synaptic plasticity | **Fig. 7, Supplementary Fig. 7** |

Under this approximation, *r*_on_ and *r*_off_ would be the same for every neuron, therefore we have

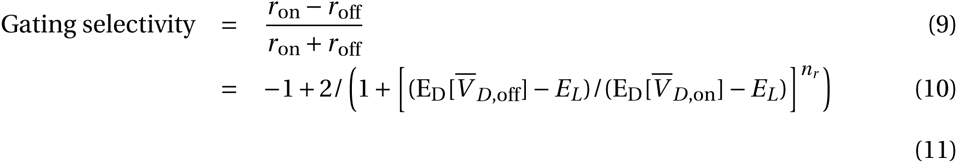

Since dendritic voltage is determined by the total excitatory and inhibitory conductance received,

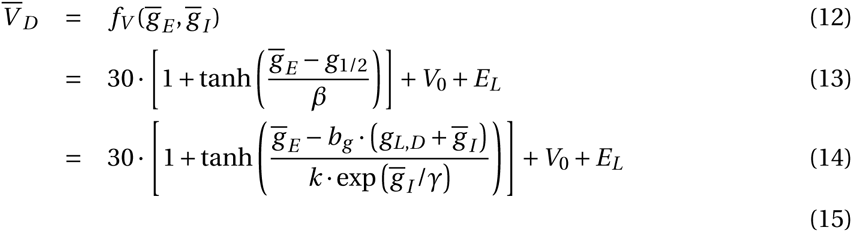

Remember that for each pathway, we assume that the excitatory input conductance is a deterministic function of the inhibitory conductance received when the corresponding gate is open.

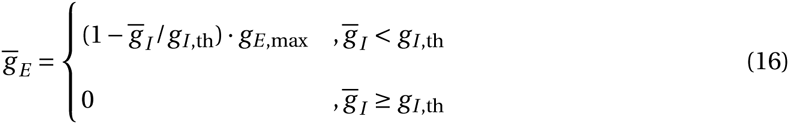

Denote this rectified linear function as 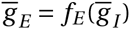. For convenience consider two pathways, the inhibitory conductance for gate 1 is 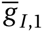 and for gate 2 is 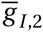. And excitatory conductance for pathway 1 and 2 are 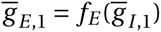 and 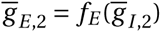 respectively. Then

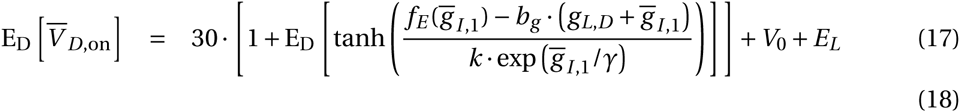

and

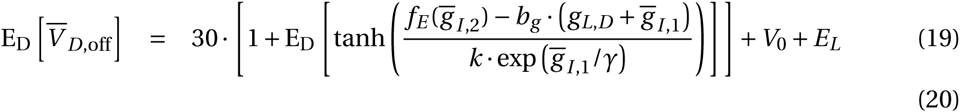

We assumed that each dendrite is targeted strictly by *N*_SOM→dend_ SOM neurons, and since we are keeping the total amount of inhibition *G*_SOM→dend_ received by each dendrite fixed, the time-averaged conductance of each connection is*G*_SOM→dend_/*N*_SOM→dend_. We also assumed that each SOM neuron gets suppressed with probability 1-*p*. Then the number of non-suppressed SOM neurons targeting each dendrite *n*_SOM→dend_ follows a binomial distribution

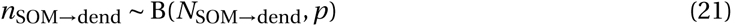

And

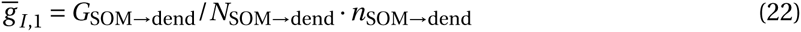

Therefore *N*_SOM→dend_ determines the distribution for 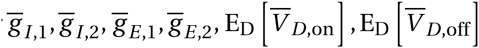, and finally the gating selectivity. In summary, in the limit of a large number of dendrites, we have shown that gating selectivity only depends on the parameter *N*_SOM→dend_.

### 3.2 Gating selectivity strictly improves with somatic inhibition

Denote the f-I response function of the somatic compartment as *ƒ*(·), and assume the dendritic input current to the soma is *I*_on_ and *I*_off_ when the gate is open or closed respectively. Also denote the somatic inhibitory current as *I*_PV_. For convenience, assume *I*_PV_ > 0, so the outputs of the pyramidal neuron are

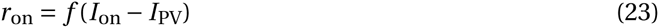

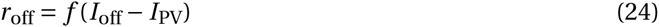

respectively. We consider only the case when *r*_on_, *r*_off_ > 0, which means input stimuli have a net excitatory effect. Also we have *I*_PV_ < *I*_off_. Since *r*_on_, *r*_off_ are baseline corrected, we should have *ƒ*(0) = 0. Here we derive the necessary and sufficient condition for gating selectivity

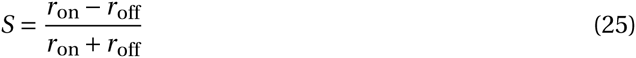

to strictly increase with *I*_PV_.

We have

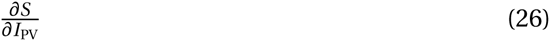

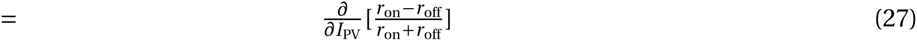

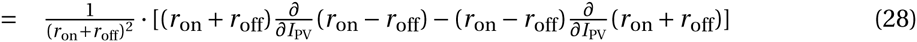

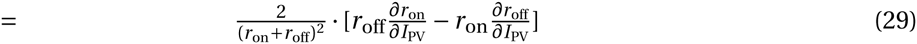

So

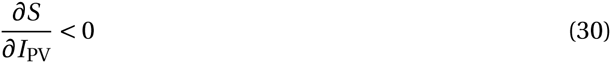

is equivalent to

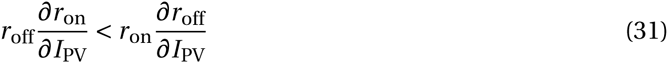

In a few more steps, wecan easily derive that the necessary and sufficient condition for gating selectivity to improve with somatic inhibition is that

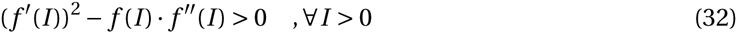

where 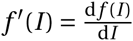.

We can easily see that for any power law function *ƒ*(*I*) = *aI*^*b*^,

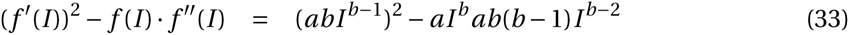

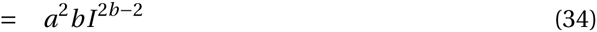

is strictly larger than 0, as long as *b* > 0.

